# Restriction-modification systems have shaped the evolution and distribution of plasmids across bacteria

**DOI:** 10.1101/2022.12.15.520556

**Authors:** Liam P. Shaw, Eduardo P. C. Rocha, R. Craig MacLean

## Abstract

Many novel traits such as antibiotic resistance are spread by plasmids between species. Yet plasmids have different host ranges. Restriction-modification systems (R-M systems) are by far the most abundant bacterial defense system and therefore represent one of the key barriers to plasmid spread. However, their effect on plasmid evolution and host range has been neglected. Here we analyse the avoidance of targets of the most abundant R-M systems (Type II) for complete genomes and plasmids across bacterial diversity. For the most common target length (6 bp) we show that target avoidance is strongly correlated with the taxonomic distribution of R-M systems and is greater in plasmid genes than core genes. We find stronger avoidance of R-M targets in plasmids which are smaller and have a broader host range. Our results suggest two different evolutionary strategies for plasmids: small plasmids primarily adapt to R-M systems by tuning their sequence composition, and large plasmids primarily adapt through the carriage of additional genes protecting from restriction. Our work provides systematic evidence that R-M systems are important barriers to plasmid transfer and have left their mark on plasmids over long evolutionary time.

## Introduction

When DNA enters a bacterial cell from the world outside, it is an unknown quantity. If transcribed into RNA then translated into protein by the cell’s own molecular machinery, the consequences may be beneficial – the survival of an unanticipated stress through the acquisition of new genes – but they may also be disastrous. Mobile genetic elements (MGEs) such as lytic phage attempt to hijack cellular machinery to their own advantage: the transcription of phage DNA leads to copies of phage being produced at the expense of the bacterial host, followed by lysis and cell death. For this reason, bacteria have evolved many ‘defense systems’ which offer protection against external DNA. Defense systems impair or block infection by MGEs. Their evolution is closely linked to MGEs (1) and they help to shape routes of gene flow between bacteria (2). The majority of prokaryotic genomes contain at least one R-M system (83%) making them by far the most abundant defense systems – over twice as abundant as CRISPR-Cas (3). R-M systems recognise specific DNA motifs and are grouped into four broad types I-IV (4).

Within R-M systems, Type II are the most abundant, present in 39.2% of bacterial genomes (3) with a mean of ∼0.5 systems per genome (5). Type II R-M systems consist of two enzyme activities: a restriction endonuclease (REase) which cuts double-stranded DNA (dsDNA) at targets and a methyltransferase (MTase) which modifies targets to protect them from cleavage. These enzymes are typically encoded by separate genes located close together in the genome. The targets of restriction are short sequences of 4-8 bases which are usually palindromic i.e. they are equal to their own reverse complement (6) due of the symmetrical subunits of the protein multimers that recognize the target (7,8). Any occurrences of the restriction target in the cell’s own DNA should be protected from restriction by the methyltransferase. In contrast, DNA originating from a different species or strain that does not have the same R-M system should lack this methylation at target sites, and will be cleaved by the restriction endonuclease when the DNA enters the cell.

R-M systems are the most-studied class of defense systems and have been heavily investigated since their discovery in the 1960s (9,10). Their widespread prevalence across bacteria suggests they provide an important defense against MGEs, which implies a strong selective pressure on MGEs to evade their targeting. Work on the first sequenced phage genomes in the 1980s showed evidence of selection against restriction targets (11) which was backed up by subsequent research (12–15). By providing an innate or ‘first-line’ immunity, R-M systems can impair incoming MGEs prior to the activation of other ‘second-line’ defense systems. They are compatible with CRISPR-Cas (16) and restriction endonuclease cleavage of viral DNA can stimulate the subsequent adaptive CRISPR response (17).

As well as functioning as defense systems, R-M systems can also be viewed as selfish elements that serve to propagate themselves. When the MTase decays more quickly than the REase, a Type II R-M system can function as an addiction system to ensure its own persistence (18,19), similar to toxin-antitoxin systems (20). This addictive quality may contribute to their occasional occurrence on MGEs such as plasmids: around 10.5% of plasmids carry R-M systems (5) and experiments have shown R-M system carriage can lead to increased plasmid stability in cells (19).

Despite the different interpretations of the evolutionary role of R-M systems, it is clear that they shape pathways of gene flow between populations. In line with this, bacteria possessing cognate R-M systems have higher rates of horizontal gene tranfsfer between them (22). One major route of this gene flow is plasmid transfer. Plasmids are vehicles for novel traits that are beneficial across species (23) including antibiotic resistance (24). However, plasmid transfer is constrained by taxonomic boundaries (25,26). The host range of a plasmid is defined as the range of different bacteria it can infect, with plasmids traditionally divided into ‘narrow’ or ‘broad’ host range. It has been suggested that plasmids with narrower host ranges tend to have a similar sequence composition to their host chromosomes (27). This would be expected due to amelioration – the tendency of horizontally transferred genes to increasingly resemble their recipient genome in sequence composition over time due to mutational biases (28) – but could also result from adaptation to the host defense systems.

More recent large-scale analyses of plasmids have quantified host range by grouping similar plasmids into clusters (25,26). These studies suggest many plasmids have a limited observed host range: considering only plasmid taxonomic units (PTUs) containing at least four plasmids, 45% are observed only in a single species (26). As barriers to the spread of dsDNA MGEs, R-M systems contribute to shaping the possible routes of plasmid transfer (21). Yet existing studies of R-M systems and plasmids are experimental and mostly limited to transfer within a single species – for example, in *Helicobacter pylori* (29) or *Enterococcus faecalis* (30).

Over fifty years ago Arber and Linn speculated that because ‘transferable plasmids have a fair chance of alternating rather frequently among hosts of various specificity…[we should] expect that with relatively small DNA molecules many original sites for the specificities of the most common hosts have been lost’ (7). Yet despite the detailed characterisation of R-M systems compared to other defense systems (31) and their ubiquity across bacteria, we still do not know whether this hypothesis holds true across plasmids. As such, we lack a systematic understanding of the role of R-M systems in shaping plasmid transfer routes across known bacterial diversity.

Here we investigate the avoidance of Type II restriction targets in plasmids, using a dataset of 8,552 complete genomes from 72 species containing 21,814 plasmids, as well as a separate dataset of plasmids with information on host range (26). Our results both confirm that avoidance of restriction targets is a general feature of bacterial genes and suggest that it may be greater in plasmids for 6-bp targets. By analysing the taxonomic distribution of Type II R-M systems and plasmids together, we show that avoidance patterns are associated with a plasmid’s size and host range: small and broad host range plasmids show greater avoidance of R-M targets. Our findings suggest that Type II R-M systems are important drivers of plasmid evolution and shape routes of plasmid transfer in bacterial populations.

## Materials and Methods

### Predicting Type II R-M systems

Our analysis approach requires a presence/absence database of R-M systems targeting particular motifs across different species of bacteria. We therefore first developed a pipeline ‘rmsFinder’ to detect Type II R-M systems and then predict their target motifs: (https://github.com/liampshaw/rmsFinder). Previous work (Oliveira, Touchon, and Rocha 2016) determined protein similarity thresholds (% amino acid identity) above which enzymes are likely to have the same target specificity: 50% for restriction endonucleases (REases) and 55% for methyltransferases (MTases). We used these as default values to define predicted targets. rmsFinder uses previously published hidden Markov models (HMMs) from either Oliveira, Touchon, and Rocha (2016) (--hmm oliveira) or Tesson et al. (2022) (--hmm tesson) to find putative Type II REases and MTases in a proteome. Here, we report results using the ‘tesson’ HMMs (those from DefenseFinder). rmsFinder then compares these putative enzymes to those enzymes in REBASE (31) which have known or previously predicted targets.

In rmsFinder, we define the presence of a Type II R-M system as the presence of an MTase and REase with a shared predicted target within 4 genes of each other (i.e. separated by at most 3 intermediate genes). rmsFinder returns both a list of possible hits to MTases and REases as well as this final prediction of Type II R-M systems with a known target. This final level of prediction can operate using different subsets of REBASE enzymes at decreasing levels of stringency:

- ‘gold’ - REBASE ‘gold standard’ proteins for which the biochemical function has been experimentally characterized and the nucleotide sequence coding for the exact protein is known.
- ‘nonputative’ - REBASE proteins that are known to have biochemical function (i.e. excluding proteins predicted bioinformatically by REBASE based on protein similarity).
- ‘all’ - all REBASE proteins, including putative protein sequences predicted bioinformatically by REBASE based on similarity to existing proteins.

Results presented in this manuscript are from the ‘all’ mode of rmsFinder using REBASE v110 (downloaded 19 October 2021). We use the proteins defined within REBASE as Type II REases or MTases. We investigated the possibility of predicting the targets of Type IIG systems where the restriction and methylation functions are encoded in a single enzyme, but found that this was not reliable (data not shown) and so restricted our analysis only to Type II systems where the REase and MTase are separate enzymes.

### Overall pipeline

We developed a pipeline to run rmsFinder on downloaded genomes from a different bacterial species to create a database of putative R-M systems with predicted targets (Supplementary Data 2) and also to compute exceptionality scores for all possible k-mers. The github repository for this paper contains analysis scripts (https://github.com/liampshaw/R-M-and-plasmids); here we describe the overall approach.

### Species genomes

We downloaded genomes for all n=104 species with >25 complete genomes in NCBI RefSeq (as of 20 January 2022) then filtered them for quality with PanACoTA v1.3.1 (32) with the ‘prepare’ subcommand (--norefseq, otherwise default parameter, meaning retained genomes have a maximum L90 of 100 and a maximum of 999 contigs). After filtering, n=72 species had >25 complete genomes (8,552 genomes in total; ‘RefSeq:>25’ dataset; for list of accessions see Supplementary Data 3). For each species, we used PanACoTA v1.3.1 to annotate genes and then perform a pangenome analysis. We defined a gene family as ‘core’ if >99% of genomes had exactly one member (corepers subcommand of PanACoTA with ‘-t 0.99 -X’). This is a more relaxed definition than a strict core genome where all genomes are required to have exactly one copy of each core gene; such a definition can produce reduced core genomes when using public genomes, because an error in any single assembled genome can remove a gene from the core genome. After annotating to find CDSs, we split each RefSeq genome into three gene components: core genes on the chromosome (‘core’), non-core genes on the chromosome (‘non-core’), and genes on other replicons (‘plasmid’). Three species in our dataset contained secondary chromosomes: *Burkholderia pseudomallei* (81/91 isolates), *Vibrio cholerae* (57/70) and *Vibrio parahaemolyticus* (43/43). For the purposes of our analysis, we treated genes on these secondary chromosomes as ‘plasmid’ genes (excluding them did not change our conclusions). We analysed target avoidance both for the entire genome and for each pangenome component separately.

### Plasmid genomes

We downloaded the dataset of n=10,634 plasmids previously analysed by Redondo-Salvo et al. (26). We used their existing classification of these plasmids into plasmid taxonomic units (PTUs). Redondo-Salvo et al. define the host range of a PTU from I-VI based on its observed distribution across taxonomic levels, from narrow (I: within-species) to broad (VI: within-phylum) (see Supp. Dataset 2 of that paper). We filtered the plasmids to n=4,000 plasmids that were seen in species from our RefSeq:>25 dataset (using TaxName in Redondo-Salvo et al.’s Dataset S2 and disregarding extra specificity after genus and species). Host range is not strongly correlated with plasmid size (e.g. for k=6 linear model dataset, Spearman’s *ρ*=0.046, p=0.10), so we include both of these variables.

### R-M target distribution

We ran rmsFinder on the 8,552 filtered genomes in our dataset of 72 species. We detected 8,616 putative R-M systems with a predicted target motif, with 2,592 genomes containing at least one R-M system (30.3%). Some putative R-M systems ‘overlapped’ e.g. the same REase could be included in multiple putative systems if there were multiple MTases in close proximity. To avoid overcounting these systems, we can consider the unique targets recognised per genome. Of the R-M-containing genomes, 1,875/2,592 (72.3%) had R-M system(s) recognising just one motif (range: 0-18 putative R-M systems; *Helicobacter pylori* genomes accounted for all those with >9 R-M systems). Six species contained no predicted R-M systems (*Bacillus anthracis, Chlamydia trachomatis, Corynebacterium pseudotuberculosis, Limosilactobacillus reuteri, Mycobacterium tuberculosis, Piscirickettsia salmonis*). R-M systems targeted 104 known REBASE motifs corresponding to 341 unambiguous sequences (hereafter: ‘targets’) of which the majority were between 4 and 6 bases long (Table 1). Where a motif contained ambiguity codes (e.g. ATNNAT) we include all possibilities as independent targets i.e. with equal weighting compared to unambiguous targets. Out of the 99 motifs of 4-6 bases, 30 were targeted by only a single species. On average, a given REBASE motif was targeted by systems found in a median of 3 species (range: 1-28) and 20 genomes (range: 1-615).

**Table 1:** Detected Type II R-M targets across the dataset of 8,552 genomes. * Number of genomes with at least 1 R-M system targeting a target of length *k*.

| Length ($k$) | REBASE motifs | $k$-mer targets | Palindromes | Genomes* | Species |
| --- | --- | --- | --- | --- | --- |
| 4 | 11 | 11 | 10 of 12 | 691 | 33 |
| 5 | 30 | 59 | - | 1266 | 61 |
| 6 | 58 | 179 | 45 of 64 | 1446 | 55 |
| 7 | 4 | 28 | - | 61 | 3 |
| 9 | 1 | 64 | - | 5 | 4 |

We then aggregated these results by species into a binary presence/absence matrix of species against k-mers for k=4,5,6 (Supplementary Data 4-6). In this matrix entries are either 1 (denoting that a functional R-M system targets the k-mer), or 0 (denoting that no R-M system was observed in the dataset targeting the k-mer). We took complete taxonomic classifications for the 72 filtered species from SILVA (33) (Table S1). For a given species, we can therefore define the set of motifs that are targeted by R-M systems observed within-species, within-genus, within-family etc. up to the order of phylum. This ‘taxonomic dictionary’ allows us to explore how the distribution of R-M systems is linked to avoidance of their associated targets in bacterial genomes and plasmids.

### Calculating target avoidance

Sequence composition strongly affects the number of times a short motif appears in a stretch of DNA. For example, one would expect few occurrences of GGCC in an AT-rich genome. We therefore used R’MES (34) to calculate an exceptionality score for all *k*-mers (*k=*4,5,6). R’MES controls for sequence composition by using a Markov chain model to calculate the expected occurrences of a word *W* of length *k* using the observed occurences of shorter words. This gives a null expectation which can be compared with the actual occurences of *W* to produce an exceptionality Z-score. For our analyses, we used R’MES v3.1.0 (https://forgemia.inra.fr/sophie.schbath/rmes) and the maximal model of order *m*=*k*-2 which uses the observed occurrences of all words with lengths ≤ *k-1* (35). The use of a maximal Markov model has the advantage that when a *k*-mer is observed significantly less than expected under the null model, this is a strong sign of selection against the word itself, rather than against the substrings it contains. Where a *k*-mer has zero observed occurrences and zero expected occurrences, its score as calculated by R’MES is defined as zero. Using the taxonomic dictionary of the presence of systems targeting particular R-M targets we then calculated the median exceptionality score for defined groups of targets for each species. For example: assume that for a given species S_a_, we detect R-M systems which target k_1_, k_2_ and k_3_. A different species S_b_ within the same genus has R-M systems targeting k_1_, k_4_ and k_5_. The within-species R-M targets of S_a_ are {k_1_, k_2_. k_3_} and the within-genus targets are {k_1_, k_2_, k_3_, k_4_, k_5_}. This logic extends up the taxonomic hierarchy, up through family, order, class, phylum and finally to kingdom, the set of targets includes all k-mers targeted by any R-M system detected within our dataset. We use only the presence of an R-M system and do not use any prevalence information.

### Controlling for sequence length

The statistical power to detect significant deviation in the abundance of motifs compared to expectation increases with sequence size. To control for differences in length between genome components, we ran analyses on both whole sequences and also subsampled sequences down to fixed lengths (2.5, 5, 10, 50, and 100 kbp) to verify that observed patterns held for fixed lengths of sequence.

### Modelling palindrome avoidance controlling for phylogeny

Genome composition is correlated with phylogeny and public databases are unevenly sampled, making overall findings about ‘average’ effects from comparative studies potentially misleading. Phylogenetically controlled analyses are required to draw reliable conclusions (36,37). We modelled the difference in R-M target avoidance between plasmid genes and core genes on the chromosome at a within-isolate level, subsampling to 10kbp; n=4,553 genomes across 60 species with at least 10kbp in each of the three pangenome components components (‘core’, ‘non-core’, and ‘plasmid’). Differences between plasmids and chromosomes can be biased by the phylogenetic structure of bacteria. To account for this, we followed the methodology of Dewar et al. (38). For the species phylogeny, we constructed a 16S rRNA gene phylogeny as follows. First, we downloaded all available nucleotide sequences from NCBI’s Bacterial 16S Ribosomal RNA RefSeq Targeted Loci Project (PRJNA33175). We then searched for our species, picked one sequence per species (the first one in the combined fasta file), aligned these sequences with mafft v7.490 (default options) (39), built a tree with FastTree v2.1 (-gtr model) (40), and then midpoint-rooted the tree before using it in modelling. This phylogeny is provided in supplementary material (Supplementary Data 7 and Fig. S1). The phylogeny can be converted into an inverse matrix of relatedness between species, which can then be used to incorporate phylogenetic structure into the random effect of species. We used MCMCglmm v2.34 (41) to model mean ranks of avoidance. The model contains pangenome component (core/non-core/plasmid) as a fixed effect and two random effects: species (with underlying phylogenetic structure) and number of genomes of a species.

### Software

All python and R code is available on github. Bioinformatic analysis of genomes and plasmids was carried out using the Biomedical Research Computing (BMRC) facility at the University of Oxford. We conducted downstream analyses in R v4.1.2 and RStudio v2022.07.2 using the following R packages: ape v5.6-1, cowplot v1.1.1, dplyr v1.1.1, formatR v1.14, ggbeeswarm v0.6.0, ggplot2 v3.4.1, ggrepel v0.9.3, ggridges v0.5.4, ggtree v3.2.1, MCMCglmm v2.34, phytools v1.0-3, reshape2 v1.4.4, tidyr v1.2.0, tidyverse v2.0.0.

## Results

### Avoidance of 6-bp palindromes is stronger in plasmid genes than in core genes

The pangenome of a species consists of all the gene families found in the species as a whole (43,44). MGEs are important contributors to the accessory component of the pangenome – genes which are variably present or absent in different members of the species. As defense systems, Type II R-M systems should exert a selective pressure within a pangenome for avoidance of their short targets, which are often palindromic and 4-6bp in length. Older studies have shown that both phage and bacteria avoid short palindromes (Rocha, Danchin, and Viari 2001; Sharp 1986), and one study on the 49kb backbone of the broad host range IncP-1 plasmid found an under-representation of 6-bp palindromes (45).

We hypothesised that the plasmid-borne components of the pangenome should show stronger avoidance of R-M targets than core genes carried on the chromosome. To test this hypothesis, we assembled a dataset of high-quality reference genomes for species from NCBI RefSeq (n=72 species with >25 genomes). Within each species, we separated genes into three pangenome components: genes where >99% of genomes in the species had exactly one copy (‘core’), other genes on the chromosome (‘non-core’), and all genes carried on other replicons (‘plasmid’). As an initial proxy for restriction targets, we first analysed the avoidance of short palindromes in each pangenome component for *k*=4 and *k*=6 (DNA palindromes require *k* to be even).

When testing evidence of avoidance of a specific target it is important to account for differences in sequence composition; for example, a GC-rich sequence should *a priori* contain fewer occurrences of an AT-rich target. To do so, we used a maximal Markov model to calculate an exceptionality score for each *k*-mer (35). Positive values of the exceptionality score for a *k*-mer (>0) indicate evidence of over-representation and negative values (<0) indicate avoidance (see Methods). Genes in all three pangenome components clearly avoided palindromes (exceptionality score < 0, k=6 Fig. 1a, for k=4 see Fig. S2 within Supplementary Data 1). We found that plasmid genes avoiding 6-bp palindromes significantly more on average than core and non-core chromosomal genes (p<0.001 two-sided Wilcoxon paired test, Fig. 1a). There was a significant correlation at the species level for palindrome avoidance in core and plasmid genes (Fig. 1b), as would be expected based on previous observations of similarities in sequence composition between plasmids and their hosts (25) and work on amelioration of genes acquired by horizontal transfer (28).

**Fig. 1.**
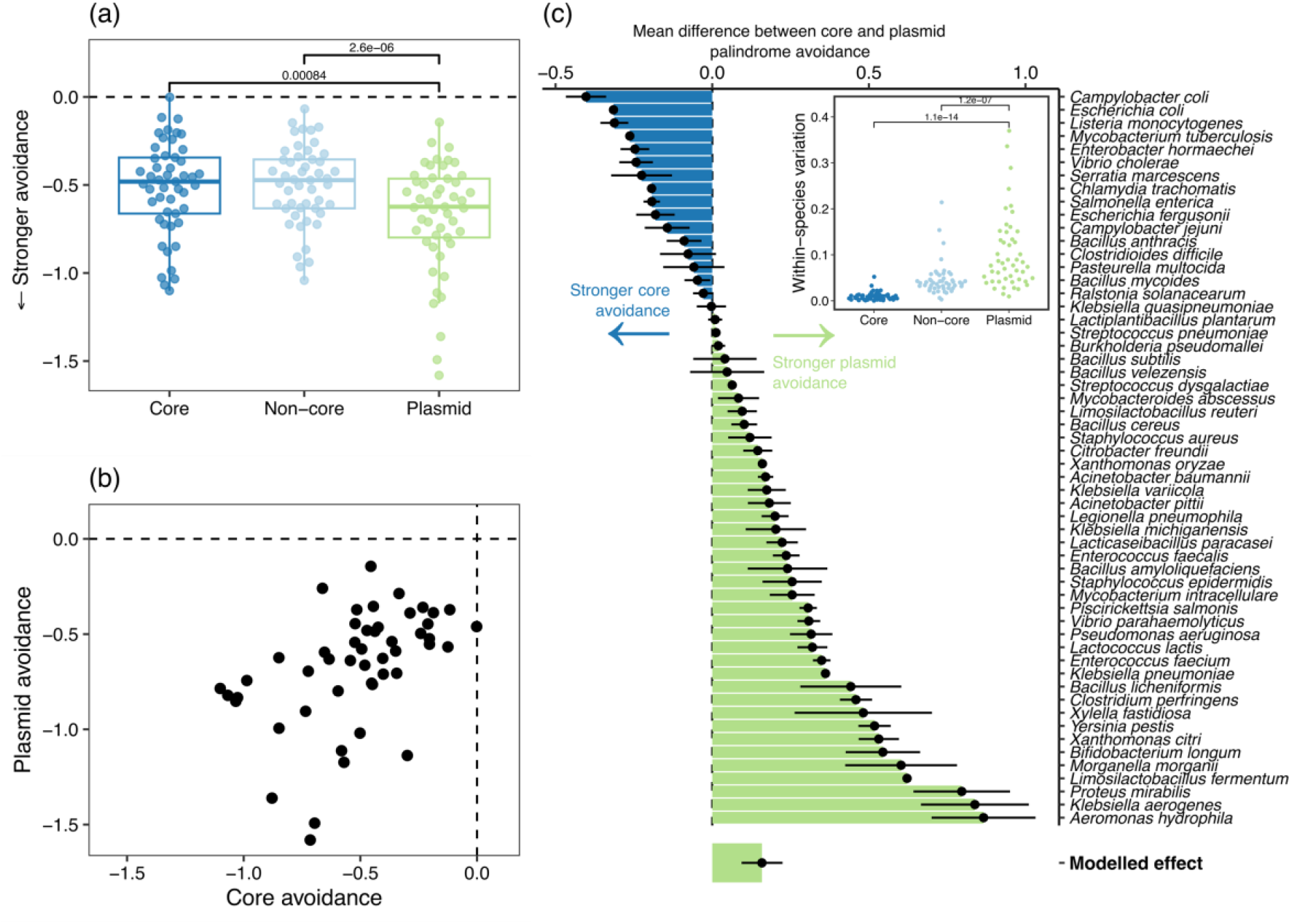
Avoidance of short palindromes (k=6) is stronger but more variable in plasmids. (a) Significantly greater avoidance of 6-bp palindromes in plasmid genes compared to core and non-core chromosomal genes (p<0.001, two-sided Wilcoxon paired test). (b) Mean avoidance is strongly structured by species, with a strong correlation between avoidance in core and plasmid genes (Spearman’s ρ=0.55, p<0.001). (c) Relative palindrome avoidance for species for core vs. plasmid genes (>0 denotes greater avoidance in plasmid genes). Points are mean, error bars show standard error. The modelled effect was computed using a phylogenetically-controlled GLMM (see Methods). Data shown are mean avoidance scores of 6-bp palindromes (4^3^=64) calculated with R’MES after pangenome construction then subsampling each per-isolate pangenome compoment to 50kbp i.e. only genomes with at least 50kbp are included (3,912 isolate genomes across 44 species). The inset panel shows within-species variation in mean palindrome avoidance score for each pangenome component. Only species with at least 3 genomes meeting these criteria are shown. For 4-bp palindromes, there was no significant difference between plasmid and core genes (Fig. S2) and mean avoidance was uncorrelated with 6-bp palindrome avoidance (Spearman’s ρ=0.005, Fig. S3). Notably, in a rare previous study, Wilkins et al. (45) found that 4-bp palindromes were not strongly avoided in the IncP-1 backbone and suggested that R-M systems with 6-bp targets were a stronger selective pressure, in line with our findings here.

However, despite this correlation we found a difference in palindrome avoidance between plasmid genes and core genes. For 6-bp palindromes, plasmid genes showed an overall greater avoidance than core genes despite variability between species (Fig. 1c; R^2^=7.1%, Table S2b). This modelling measures the effect after accounting for any phylogenetic signal and number of sampled genomes by using generalized linear mixed models (GLMMs) (41), (see Methods and Table S2). Notably, variation in palindrome avoidance was much greater in plasmid genes than core genes (Fig. 1c, inset panel) consistent with the expectation that plasmids seen within a species may have diverse evolutionary histories. This greater variability suggests the importance of considering differences between individual plasmids.

### The taxonomic distribution of Type II R-M systems correlates with target avoidance

Our genomic dataset spanned a wide range of bacterial diversity (Fig. S3). We hypothesised that selective pressure for avoidance of a target should correlate with the frequency of encounter with an R-M system targeting it. Reliable prediction of targets for novel sequences is only possible for Type II R-M systems where restriction and methylation are carried out by different enzymes (22) (see Methods). We developed a pipeline (‘rmsFinder’) to predict both the presence and targets of Type II R-M systems in our dataset using the curated REBASE database of known R-M enzymes. We produced a presence-absence matrix of k-mers targeted by Type II R-M systems across species in our dataset: when we detected a system with a target *t* in a genome from species *s*, we classed *t* as a within-species restriction target of *s*. In turn, we used this presence-absence matrix to produce a taxonomic dictionary of targets for each species (Fig. 2a-c), ranging from within-species to within-phylum targeting based on the detected presence of R-M systems across our dataset. We detected 8,616 putative R-M systems where we could confidently predict their target and 2,592 genomes contained at least one R-M system (30.3%). Of these putative systems, 8,100 (94.0%) were carried on the chromosome. R-M systems targeted 104 known REBASE motifs. Accounting for ambiguous bases, R-M systems targeted 341 specific *k*-mers, the majority of which (52.5%) were 6-bp targets (Table 1). Since motifs of *k*=7 and 9 were not prevalent (only observed in 66 genomes) we analysed targets for *k*=4,5,6 (99/104 motifs; Table 1) across our pangenome dataset. Type II R-M systems for these targets showed a highly variable presence/absence distribution across species (Fig. S4-S6 for different *k*).

**Fig. 2.**
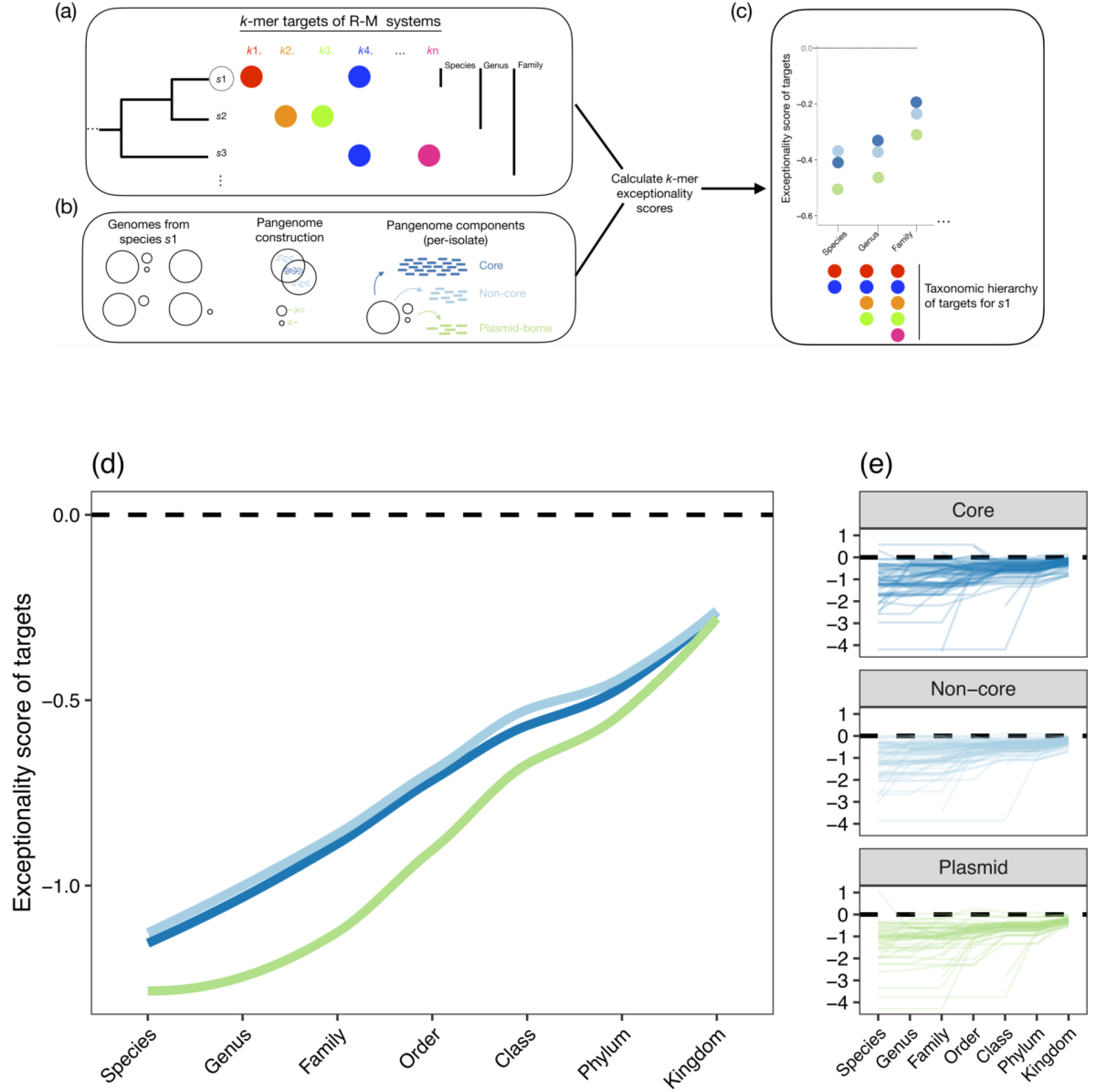
The taxonomic distribution of R-M systems correlates with avoidance of their targets. (a-c) Methodological approach to connect Type II R-M system distribution to target avoidance: (a) We search for Type II R-M systems in n=8,552 genomes from 72 species, detecting complete systems with confident prediction of targets them in 2,740 genomes (Table 1). From these hits, we created a taxonomic hierarchy of their targets across a set of species. (b) We constru ct a pangenome for each species in our dataset, then separate each individual isolate into genes in three pangenome components: core, non-core and plasmid. (c) We subsample pangenome components to a fixed size and use R’MES to calculate exceptionality scores for fixed-length *k*-mers for *k*=4,5,6 for each species, using the taxonomic hierarchy of R-M targets to correlate exceptionality scores with R-M distribution. (d-e) Exceptionality scores for 6-mers by pangenome component as a function of the taxonomic hierarchy of R-M targets: (d) averaged over all species and (e) for individual species. Subsampling is to 50kbp for each within-isolate pangenome component. Other subsampling lengths show the same pattern (see github repository).

For all pangenome components and all *k*, avoidance of targets was strongly correlated with the taxonomic distribution of the associated R-M systems (*k*=6 Fig. 2d-e; k=4 Fig. S7 and *k*=5 Fig. S8). Species pangenomes have the greatest avoidance of targets of the R-M systems found within that species. Core and non-core chromosomal genes had highly similar avoidance patterns. Selective pressure from R-M systems has imposed selection for plasmids to avoid R-M targets, and the strength of this avoidance seems to be proportional to their frequency of encounter, with the same qualitative pattern as core genes. This is consistent with the hypothesis that R-M systems are closely connected with taxonomic boundaries and plasmid host range. It is difficult to say whether R-M targets are avoided more in plasmid genes than core gene ‘on average’ across different bacteria when trying to combine different values of *k*. For *k*=6, where R-M systems contain the highest number of unique *k*-mer targets and are widely distributed, targets within the same taxonomic family were avoided more by plasmid genes at nearby taxonomic levels (species to family), with this difference decreasing at higher taxonomic orders (class, phylum) to no difference when considering avoidance of all observed R-M targets within the dataset (kingdom). However, it should be noted that for *k*=4 plasmid genes had weaker avoidance than core genes (Fig. S7), perhaps in line with the lack of difference between avoidance of palindromes in core and plasmid genes (Fig. S1), and for *k*=5 there was no clear difference (Fig. S8), so we caution against generalising this result.

### The density of within-species R-M targets increases with plasmid size

It is the actual number of occurrences of a R-M target within a plasmid that determines the extent to which it will be restricted by the associated R-M system. The expected number of target occurrences increases linearly with the size of the plasmid: for a plasmid of length *L*, the probability of containing a given *k*-mer scales as ∼ L/4^k^. For a random *k*-mer, one should expect a constant mean density. However, when we examine plasmids from the most prevalent species in our genomic dataset, *Escherichia coli*, the density of R-M targets increases with plasmid size: larger plasmids have a disproportionate number of targets (Fig. 3a-b). This pattern is not as consistent across species, although the comparison between the smallest and largest plasmid sizes within a species is significant for *k*=6 (Fig. 3c-e). From another perspective, across all plasmids when considering palindromes as a proxy for R-M targets, plasmids <10kb have a lower mean palindrome density than those >100kb for both k=4 and k=6 (p<0.001 WIlcoxon test, Fig. S9). Particularly for k=6 there is a marked increase in mean palindrome density with intermediate plasmid sizes, suggesting a gradient of densities (Fig. S10b).

**Fig. 3.**
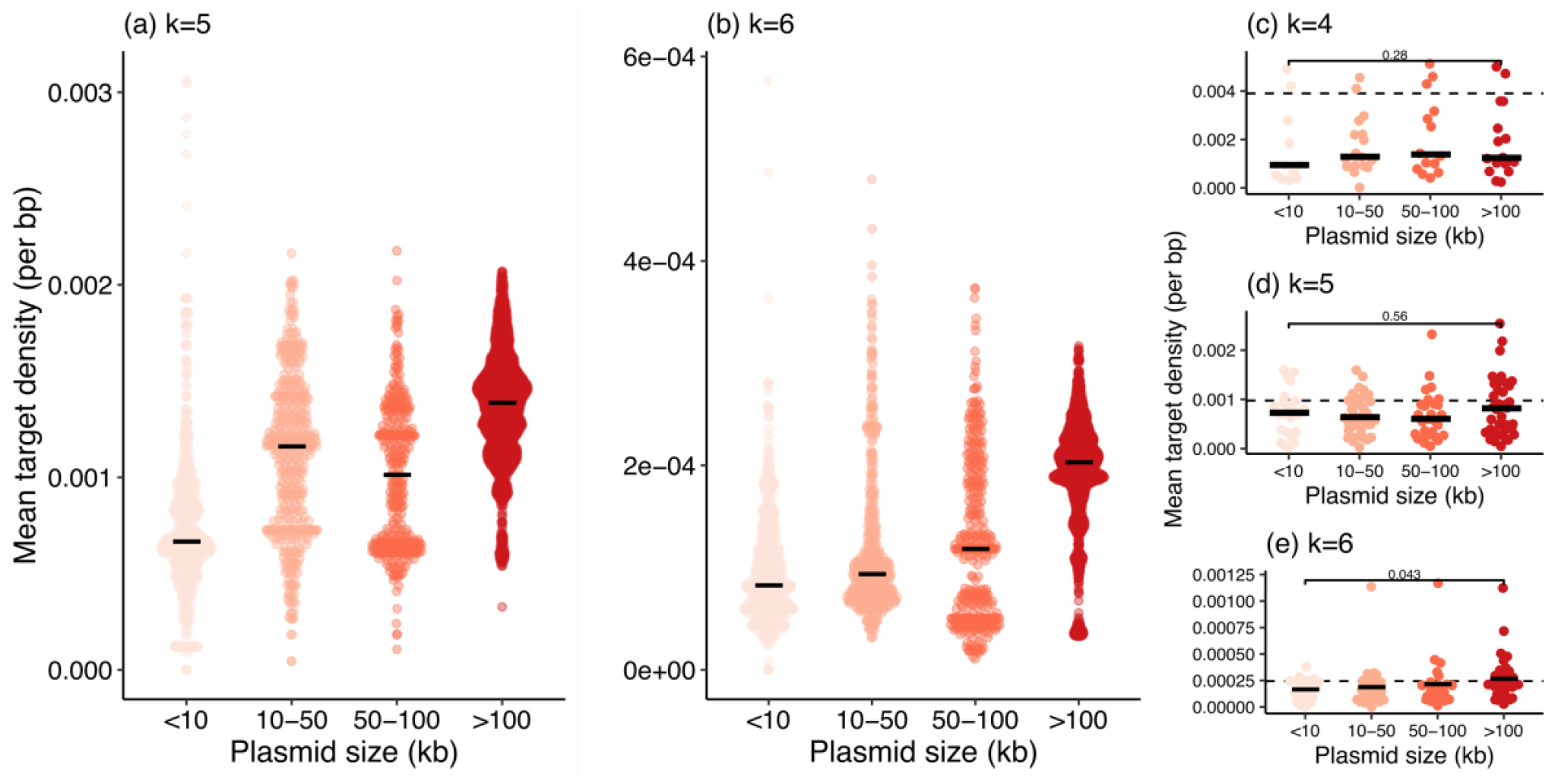
Larger plasmids have a higher density of the targets of within-species R-M systems. (a-b) Results for the best-sampled species in our genomic dataset, *Escherichia coli*, for the mean density of within-species R-M targets of length (a) *k*=5 (4 targets) and (b) *k*=6 (33 targets). Each point is the mean density of targets within a single plasmid (no deduplication), black lines show median for each category. (c-e) Results for at a per-species level for different values of *k*. Species without R-M systems with targets of length *k* are omitted. Each point represents the median of the mean densities of within-species R-M targets (both forward and reverse strand) for plasmids in that species, including only size/species combinations with >5 plasmids. Dashed lines shows the expected) density of a random *k*-mer in a random sequence d=4^-k^ (we normalise the combined forward and reverse counts by a factor of 2). The comparison between the largest (>100kb) and smallest (<10kb) plasmid category is significant (p<0.05) for *k*=6 but not *k*=4,5.

From an evolutionary perspective, a lower density of R-M targets in smaller plasmids is consistent with the fact that selective pressure from R-M systems acts at the whole-plasmid level. The efficiency of R-M systems in restricting sequences should increase with target frequency, although some systems can restrict sequences with only a single target and others require two targets to function (46,47). R-M systems thus exert a selective pressure for target depletion: without other avoidance mechanisms, to avoid restriction a plasmid must lose the restriction targets from its sequence. The number of targets, and thus the number of mutations required to lose them, increases with plasmid length.

By way of an example, consider the case of a target of length *k*=6. Each extra 5kb of sequence will, on average, add ∼1 more occurrence of the target (4^6^=4,096). At one extreme, for a small 5kb plasmid, losing its only copy of the target requires only one mutation. This mutation will carry a large fitness advantage. However, larger plasmids will require many more mutations to become target-free: a 100kb plasmid will contain ∼20 copies. While the final target-free sequence will have a large fitness advantage relative to its initial state, it must be reached gradually. Each mutational step will likely have only a weakly positive advantage compared to the previous step. Therefore, the larger a plasmid gets, the less evolutionarily accessible the mutational route to evade R-M systems becomes. The clear increase we find in the density of R-M targets with plasmid size across thousands of plasmids suggests that larger plasmids need other mechanisms of avoiding restriction.

The mean number of Type II R-M systems in a genome varies a great deal between species (Figure S10). Most species with plasmids have a mean of <1 R-M system per genome, but there are a few ‘R-M-rich’ species where every genome contains multiple R-M systems which can recognise different targets, notably *Neisseria gonorrhoeae* (mean 7.8 R-M systems per genome recognising unique targets, range 6-8) and *Helicobacter pylori* (11.6, range 8-18). It is striking that these species have small plasmids: the median plasmid size is <10kb (4.2kb and 8.2kb for for *H. pylori* and *N. gonorrhoeae* respectively) and no plasmid exceeds 50kb (42.9kb and 18.8kb). Furthermore, the mean number of plasmid bases in a genome is always <20kb (16.1kb and 1.5kb). It is also notable that 10 out of the 11 observed 4-bp R-M targets are targeted by R-M systems in *H. pylori* (Fig. S4); smaller targets are more challenging for large plasmids to avoid. It seems plausible that in such extreme species the abundance of R-M systems with diverse targets makes it almost impossible for large plasmids to persist.

### Plasmid host range correlates with stronger avoidance of R-M targets

Previous work by (26) clustered 10,634 plasmids based on their sequence similarity, defining 276 plasmid taxonomic units (PTUs) with at least four member plasmids (3,725 plasmids). They defined a host range for each PTU using its observed hosts, ranging from I-VI (from species to phylum). Under the hypothesis that R-M systems are a significant barrier to plasmid transfer, we would expect PTUs with a greater host range to have experienced more recent selection from a wider variety of R-M systems and therefore to have greater avoidance of R-M targets.

Using 6-bp palindromes as a proxy for Type II R-M targets, we find that host range is correlated with avoidance (Fig. 4). There is no such correlation for 4-bp palindromes (Fig. S11). Interestingly, the avoidance of 6-bp palindromes in plasmids that are not members of an assigned PTU suggests that they are most similar to PTUs with a within-species host range in terms of palindrome avoidance. Many singleton plasmids (those detected only once) are probably indeed restricted to single species, although notably there is a long tail of more negative exceptionality scores, suggesting that some may have broader host ranges and/or be more recent entrants into the pangenome of that species with more avoidance of targets of R-M systems seen outside the species. To understand the effect of plasmid size, we then modelled avoidance of R-M targets.

**Fig. 4.**
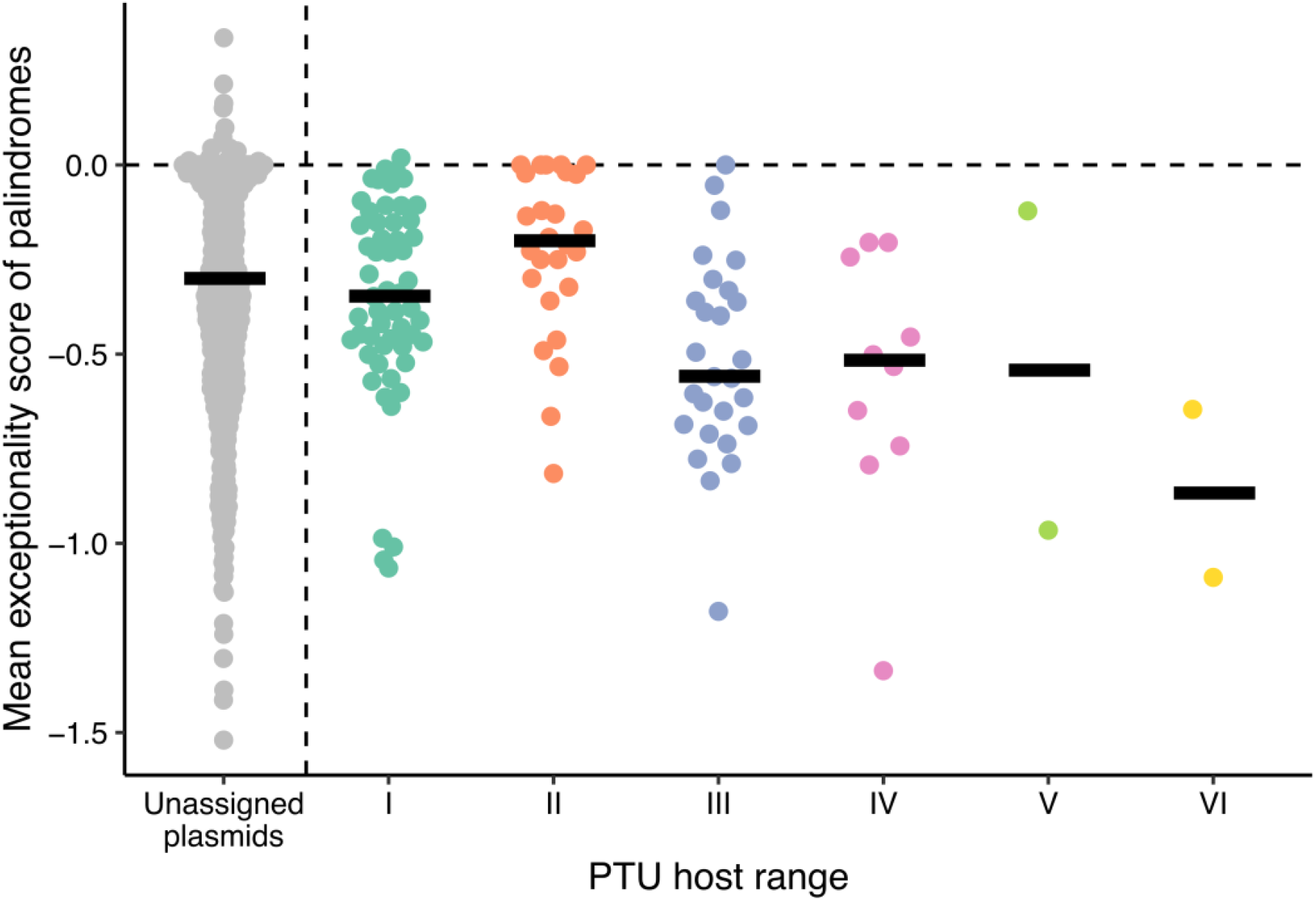
PTU host range is associated with greater avoidance of 6-bp palindromes. Avoidance of 6-bp palindromes in PTUs >10kbp correlates with PTU host range (excluding unassigned plasmids, Spearman’s *ρ*=-0.26, p=0.003). Each point is one PTU (mean exceptionality score) apart from unassigned plasmids (those not classified into a PTU) and lines show median within host ranges. There is no correlation for avoidance of 4-bp palindromes (Fig. S11).

We then modelled the avoidance of R-M targets using our taxonomic hierarchy in 4,000 PTUs seen in the same species as our dataset of complete genomes (see Methods). Linear models for exceptionality scores of 6-bp R-M targets in PTUs showed that the host range of plasmids was consistently associated with stronger avoidance of targets (Fig. 5a). In contrast, plasmid length was associated with weaker avoidance (Fig. 5b), a finding recapitulated for other values of *k*, confirming that small plasmids show greater signatures of mutational adaptation to evade R-M systems (*k*=4 Fig. S12; *k*=5 Fig. S13).

**Fig. 5.**
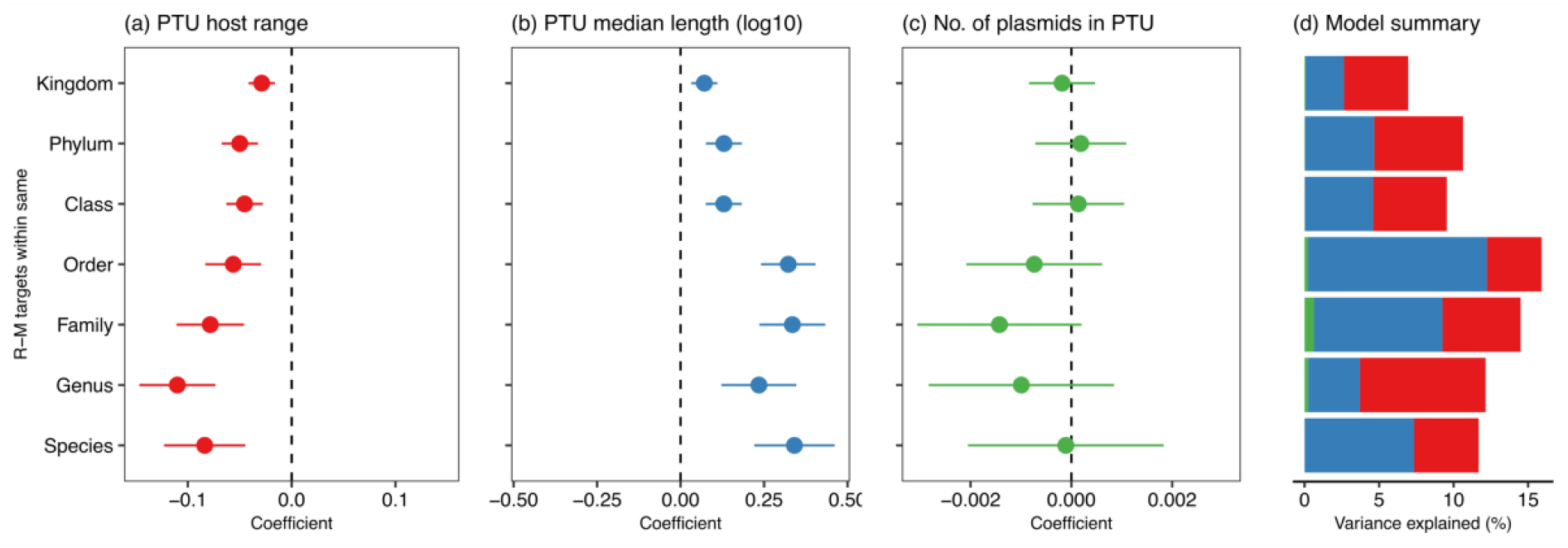
Small and broad host range PTUs have stronger avoidance of R-M targets. (a-c) Coefficients in linear models (mean estimates with standard error shown by errorbars) for the exceptionality score of R-M targets. A different model was run for each possible of level of R-M targets within the taxonomic hierarchy, from R-M targets of R-M systems within-species to within-kingdom, with three variables for each PTU: host range, median length, and number of plasmids. (a) PTU host range, converted to a numeric variable for modelling where larger values denote broader host range, is negatively associated with exceptionality score of R-M targets i.e. broader host range PTUs have stronger avoidance. (b) Median length of plasmids within PTU (log10 for modelling) is positively associated with exceptionality score of R-M targets i.e. larger plasmids have weaker avoidance. (c) Number of plasmids within the PTU has no significant effect. (d) Total variance explained by each model, with colours denoting the three different variables (red: host range, blue: length, green: number of plasmids).

The magnitude of coefficient estimates decreased in magnitude for R-M targets from progressively wider taxonomic distributions (Fig. 5a-b), consistent with avoidance patterns being signatures of plasmid adaptation to their hosts within taxonomic boundaries. The number of plasmids within a PTU did not affect its average avoidance patterns (Fig. 5c). Models explained more variance at lower taxonomic levels of R-M target distribution (Fig. 5d), with the most variance explained for PTU avoidance of R-M targets from the same order as the plasmid. Taken together, these modelling results provide strong evidence that PTUs of small size and broad-host range have greater avoidance of R-M targets. Furthermore, these effects are most noticeable for R-M targets from nearby taxonomic levels. Evading R-M targeting through mutation is an important adaptive route for small, broad host range plasmids – raising the question of how larger plasmids evade R-M systems.

### Broad host range plasmids carry more methyltransferases

The carriage of anti-restriction genes can help MGEs to evade restriction even when they carry sites recognized by the host (48). Most of these sytems remain poorly described. Yet, a well-characterised way to evade restriction is by encoding a solitary Type II MTase. Such ‘orphan’ MTases are present in many prokaryotes and likely have functions linked to genome regulation (49), but they can also provide a plasmid with effective protection against restriction against multiple R-M targets (50). Our hypothesis about the necessity of adaptation through gene carriage for large plasmids suggests that solitary MTases should be frequently carried by larger plasmids and particularly those with a broader host range.

We searched all 10,634 plasmids in the Redono-Salvo dataset for MTases and REases. Overall, only 329/10,634 plasmids (3.1%) carried at least one putative R-M system. These tended to be larger plasmids (median size 65.3kb). Considering just MTases, 1,444 plasmids carried at least one Type II MTase with a predicted target (13.6%; of which 243 carried >1 MTase), of which 789 had an MTase with a 4-6bp target (of which 173 plasmids had >1 MTase).

We looked within these plasmids for ‘orphan MTases’ where rmsFinder did not detect an associated REase recognising the same target within four genes. Larger plasmids from broad host range PTUs were more likely to carry orphan MTases (Fig. 6). Analysing at the level of PTUs and subsetting based on their size, large PTUs (>100kbp) had both a greater proportion of their members carrying MTases and a greater normalised density of MTases (Fig. S14). We modelled MTase carriage as a function of PTU median length (log10) and host range. Both size and host range were associated with MTase carriage (Table S3a). When only considering large PTUs (>100kbp; n=61 PTUs), host range was strongly associated with a greater per-base density of Type II MTases (p<0.01, adjusted R^2^=27.3%; Table S3b). Though carriage of MTases could also be linked to modulation of host chromosome gene expression, these patterns are consistent with the expected differential responses to selective pressure from R-M systems. Small plasmids rarely carry MTases but can still have a broad host range despite this because of adaptive mutations. In contrast, most large plasmids with a broad host range carry MTases.

**Figure 6.**
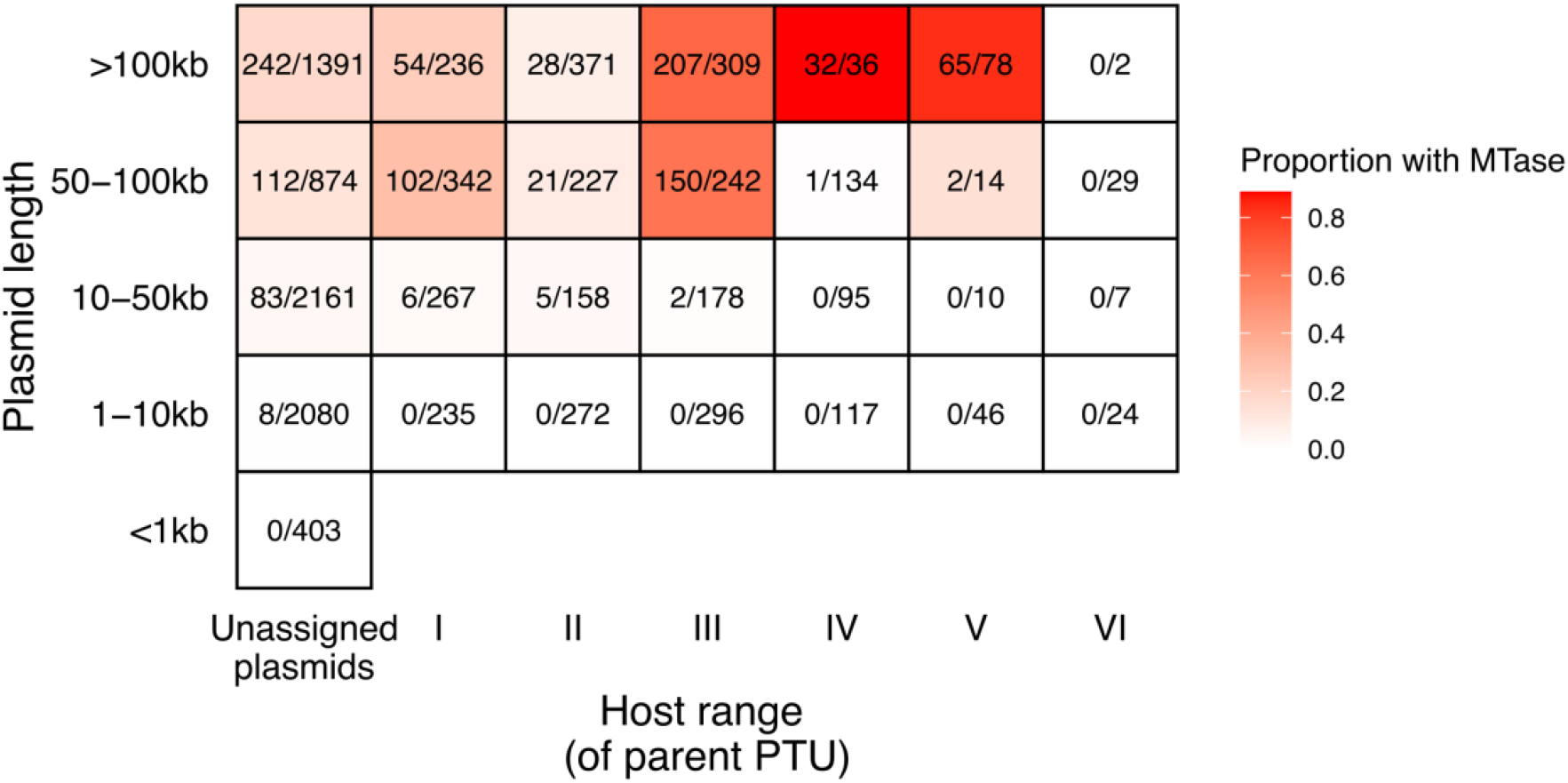
Large plasmids with a broad host range are more likely to carry ‘orphan’ MTases. Numbers show the number of plasmids in that category with at least one MTase and no putative R-M system recognising its target detected on the same plasmid, out of the total number of plasmids. We detected a putative R-M system recognising the MTase’s target for 350/1745 (20.1%) of MTase hits on Redondo-Salvo plasmids.

## Discussion

In human history, trade routes such as the Silk Road have been shaped by geography and politics; they have played an important long-term role in the movement of people, goods, and ideas. In bacterial evolution, routes of horizontal gene transfer between species have been shaped by defense systems. Here, by analysing the taxonomic distribution of the most prevalent of these defense systems – Type II R-M systems – we have shown that they have shaped the evolution and host range of plasmids. Our findings are consistent with the fifty-year-old hypothesis of Arber and Linn that small plasmids should avoid R-M targets in relation to their frequency of encounter (7).

Palindromes are important sequence features used by other DNA-binding proteins beyond R-M systems, such as the global transcriptional regulator catabolite activator protein (CAP or CRP) in *E. coli*, for which the consensus binding site is a 22-bp palindrome (51). While these other uses of palindromes may contribute to their avoidance patterns, previous studies have demonstrated convincingly that the avoidance of short palindromes (4-6bp) is a general feature of bacterial genomes that is correlated with the distribution of Type II R-M systems (12–14). While such analyses demonstrated that short palindromes are useful proxies for Type II R-M targets, they were limited in scope and not phylogenetically controlled. We have verified that these avoidance patterns persist when accounting for phylogeny across a wide range of bacteria. Although for 4-bp palindromes we found no difference in avoidance between plasmid genes and core genes, for 6-bp palindromes we found a greater avoidance in plasmid genes. Furthermore, we went beyond examining palindromes alone and showed that the taxonomic distribution of R-M systems is correlated with avoidance of their targets in all pangenome components, suggesting that R-M systems could play a role in policing species boundaries. Plasmid genes also show much greater variation, consistent with their diversity of evolutionary histories. We found that the host range of plasmid taxonomic units (PTUs) was associated with greater avoidance, suggesting that an interplay between R-M systems and plasmid host range. Models of R-M target avoidance explained the most variance for targets of systems seen within the same taxonomic order, which coincides with the observation that only 2.5% of PTUs have wider host ranges (26). We believe these findings make sense from the perspective of an evolutionary arms race between bacteria and plasmids.

We found that small plasmids had a greater avoidance of R-M targets. We argue this is consistent with the greater evolutionary ‘accessibility’ of target removal by mutation compared to large plasmids: small plasmids need fewer mutations to become target-free, and each of these mutations has a strong fitness advantage. Furthermore, smaller plasmids tend to exist at higher plasmid copy number per cell. Since multi-copy plasmids can accelerate adaptive evolution by providing a greater mutational supply (52) and avoidance of restriction is likely to be adaptive, this may contribute to an even greater depletion of restriction targets. Phage avoidance of R-M targets is greater for non-temperate phage, which have a lifestyle more dependent on horizontal transmission (15). Small multi-copy plasmids may be more ‘phage-like’ in this sense.

Plasmids have a highly bimodal size distribution: a strong peak at 5kb, very few plasmids at around 20kb, and a broad peak around 100kb (53). But their fitness costs do not seem to be correlated with their size, at least when considering resistance-carrying plasmids (54). The bimodal distribution is widely recognised, yet it presents a puzzle: if adding genes to plasmids is cheap, why do so many plasmids remain small? Plasmids are often divided into non-mobilisable, mobilisable, and conjugative plasmids. Physical considerations of horizontal gene transfer must play a role in plasmid size. First, the apparatus of conjugation and transfer machinery has a minimum size (∼10 kb), giving conjugative plasmids a minimum size. Second, there may be selection for mobilisable plasmids that are able to exploit phage mechanisms for horizontal transfer, giving these plasmids a maximum size of ∼40kb (55). As is often the case in biology, there are likely multiple contributing factors, but we suggest one that may have been overlooked is the role of R-M systems.

There are three observations that support a role for R-M systems in shaping plasmid size. First, we found that 6-bp targets were the most common Type II R-M system. The first peak in plasmid size at 5kb is the length at which the expectation of a given 6-mer is ∼1 (4^6^=4,096), making it possible to evade any 6-mer targeting system through a single mutation (for 7-mer targets, the corresponding size is ∼16.3kb). Second, the species that carry many and diverse Type II R-M systems do not have any large plasmids, suggesting that R-M systems constrain small plasmids to remain small. The small number of plasmids of ∼20kb in the size distribution of plasmids could be explained by this factor. Third, increasing plasmid size has a larger R-M-associated cost for smaller plasmids: the difference between zero and one or two copies of a target is a large one. It should be noted that some R-M systems interact with two recognition sites to cleave DNA, and more targets will probably increase the efficiency of restriction (46,47). However, once plasmids have many copies of an R-M target in their sequence, having an additional target present is unlikely to be as great a proportional burden as the first few targets. Instead, because mutational adaptation becomes increasingly difficult with plasmid size, carrying additional genes becomes the main route of adaptation: genes which allow the evasion of R-M systems (single MTases or anti-restriction enzymes) or other genes that benefit the host to increase the likelihood of vertical inheritance after breakthrough infection.

Another effect to consider is that, all else being equal, there is another reason to expect large plasmids to have a lower barrier to gene incorporation than small plasmids. If a 100kb plasmid gains a gene of ∼1kb, this represents a proportional length increase of 1%; for a 5kb plasmid, it would be an increase of 20%. If size is assumed to be broadly correlated with replicative burden, then large plasmids have a comparatively smaller barrier to incorporating new genes. Indeed, most pairs of plasmids with 95% identical relaxases exhibit less than 50% similarity in terms of their gene content (56), demonstrating that gene gain and loss in plasmids are rapid.

To summarise these lines of argument, there are good reasons to think that selective pressure from R-M systems can simultaneously drive small plasmids to become smaller and large plasmids to become larger. A similar logic applies to all defense systems targeting small DNA motifs.

Our work has limitations. Most notably, plasmid sequences are subject to a far greater range of selective pressures than we have explored here. Even considering just other defense systems alone, we have not investigated: the dual-function Type IIG enzymes with combined REase and MTase function (4), the less common but still highly prevalent Type I, III, and IV R-M systems (31) or indeed other ‘antiviral’ systems altogether (3). There is also a growing appreciation that MGEs use ‘defense’ systems as weapons of intragenomic conflict (57). Other pressures apart from defense systems may shape sequence composition: for example, there is some evidence that plasmids are AT-rich compared to chromosomes to reduce their metabolic burden (58). In restricting our analysis to Type II R-M systems we have been deliberately conservative. Although we believe our findings are consistent with their expected action against plasmids, our analysis is only a partial picture of these complex overlapping pressures. We wish to highlight that our conclusions seemed to consistently apply more to 6-bp R-M targets than other lengths (k=4,5), which may be indicative of systematic bias or possible have underlying biological reasons. For example, there are simply more possible 6-bp-targeting R-M systems and 6-mers have more freedom to change without disrupting coding sequences through synonymous changes.

In conclusion, although Type II R-M systems are usually studied through the lens of phage defense, they have also shaped plasmid evolution. The selective pressure from R-M systems manifests differently with different plasmid sizes: small plasmids primarily evade restriction by point mutations that eliminate targets from their sequences, while large plasmids with many more targets instead acquire accessory genes such as methyltransferases to protect against restriction. More generally, our work suggests that avoidance patterns in MGEs contain information on the immune pressures they have endured. At a time when many novel ‘phage defense systems’ are being discovered, analysis of avoidance patterns can elucidate how these systems may have shaped the evolution and spread of other MGEs.

## Availability

Genomes analysed are all from public databases (NCBI) and accessions are available as Supplementary Data. Analysis scripts are on github (https://github.com/liampshaw/R-M-and-plasmids) as is rmsFinder, the tool we developed to find and predict the targets of putative R-M systems (https://github.com/liampshaw/rmsFinder). The R-M- and-plasmids github repository can be used to reproduce analyses (figures and tables) using intermediate data files made available via figshare: https://doi.org/10.6084/m9.figshare.21923121.v1

## Supplementary Data

Supplementary Data are available at NAR online:

- Supplementary Data 1. All supplementary figures and tables (pdf).
- Supplementary Data 2: Putative R-M systems detected in the complete genomes analysed.
- Supplementary Data 3. List of ftps for all complete genomes analysed as part of the species whole genome dataset.
- Supplementary Data 4. Matrix of 4-mers and species with 1 indicating that an R-M system targeting that 4-mer was observed at least once in a complete genome of the species using rmsFinder
- Supplementary Data 5. Matrix of 5-mers and species with 1 indicating that an R-M system targeting that 5-mer was observed at least once in a complete genome of the species using rmsFinder.
- Supplementary Data 6. Matrix of 6-mers and species with 1 indicating that an R-M system targeting that 6-mer was observed at least once in a complete genome of the species using rmsFinder.
- Supplementary Data 7: 16S rRNA gene tree for n=72 species analysed in the genomic dataset.

## Acknowledgements

We would like to thank the curators of REBASE and its many contributors, without which this work would have been impossible. Thanks to Sergio Redondo for discussions about PTU host range, Sophie Schbath for correspondence about R’MES, Anna Dewar for discussion of phylogenetically-controlled GLMMs, and the three reviewers for their constructive comments.

