## Supplementary Data 1 for "Restriction-modification systems have shaped the evolution and distribution of plasmids across bacteria"

### Supplemental Material

For code and datasets to reproduce figures and analysis (including these supplementary figures and tables), see the paper's github repository: https://github.com/liampshaw/R-M-and-plasmids.

**Table S1. Taxonomic assignments for the 72 species in the filtered genomic dataset.** Taxonomic levels are separated by semicolons.

| Kingdom; Phylum; Class; Order; Family; Genus; Species |
|---|
| Bacteria;Proteobacteria;Gammaproteobacteria;Pseudomonadales;Moraxellaceae;Acinetobacter;Acinetobacter baumannii |
| Bacteria;Proteobacteria;Gammaproteobacteria;Pseudomonadales;Moraxellaceae;Acinetobacter;Acinetobacter pittii |
| Bacteria;Proteobacteria;Gammaproteobacteria;Enterobacterales;Aeromonadaceae;Aeromonas;Aeromonas hydrophila |
| Bacteria;Verrucomicrobiota;Verrucomicrobiae;Verrucomicrobiales;Akkermansiaceae;Akkermansia;Akkermansia muciniphila |
| Bacteria;Firmicutes;Bacilli;Bacillales;Bacillaceae;Bacillus;Bacillus amyloliquefaciens |
| Bacteria;Firmicutes;Bacilli;Bacillales;Bacillaceae;Bacillus;Bacillus anthracis |
| Bacteria;Firmicutes;Bacilli;Bacillales;Bacillaceae;Bacillus;Bacillus cereus |
| Bacteria;Firmicutes;Bacilli;Bacillales;Bacillaceae;Bacillus;Bacillus licheniformis |
| Bacteria;Firmicutes;Bacilli;Bacillales;Bacillaceae;Bacillus;Bacillus mycoides |
| Bacteria;Firmicutes;Bacilli;Bacillales;Bacillaceae;Bacillus;Bacillus subtilis |
| Bacteria;Firmicutes;Bacilli;Bacillales;Bacillaceae;Bacillus;Bacillus velezensis |
| Bacteria;Actinobacteriota;Actinobacteria;Bifidobacteriales;Bifidobacteriaceae;Bifidobacterium;Bifidobacterium breve |
| Bacteria;Actinobacteriota;Actinobacteria;Bifidobacteriales;Bifidobacteriaceae;Bifidobacterium;Bifidobacterium longum |
| Bacteria;Proteobacteria;Gammaproteobacteria;Burkholderiales;Burkholderiaceae;Burkholderia-Caballeronia-Paraburkholderia;Burkholderia pseudomallei |
| Bacteria;Campylobacterota;Campylobacteria;Campylobacterales;Campylobacteraceae;Campylobacter;Campylobacter coli |
| Bacteria;Campylobacterota;Campylobacteria;Campylobacterales;Campylobacteraceae;Campylobacter;Campylobacter jejuni |
| Bacteria;Verrucomicrobiota;Chlamydiae;Chlamydiales;Chlamydiaceae;Chlamydia;Chlamydia trachomatis |
| Bacteria;Proteobacteria;Gammaproteobacteria;Enterobacterales;Enterobacteriaceae;Citrobacter;Citrobacter freundii |
| Bacteria;Firmicutes;Clostridia;Peptostreptococcales-Tissierellales;Peptostreptococcaceae;Clostridioides;Clostridioides difficile |
| Bacteria;Firmicutes;Clostridia;Clostridiales;Clostridiaceae;Clostridium sensu stricto 1;Clostridium perfringens |
| Bacteria;Actinobacteriota;Actinobacteria;Corynebacteriales;Corynebacteriaceae;Corynebacterium;Corynebacterium pseudotuberculosis |
| Bacteria;Proteobacteria;Gammaproteobacteria;Enterobacterales;Enterobacteriaceae;Enterobacter;Enterobacter hormaechei |
| Bacteria;Firmicutes;Bacilli;Lactobacillales;Enterococcaceae;Enterococcus;Enterococcus faecalis |
| Bacteria;Firmicutes;Bacilli;Lactobacillales;Enterococcaceae;Enterococcus;Enterococcus faecium |
| Bacteria;Proteobacteria;Gammaproteobacteria;Enterobacterales;Enterobacteriaceae;Escherichia-Shigella;Escherichia coli |
| Bacteria;Proteobacteria;Gammaproteobacteria;Enterobacterales;Enterobacteriaceae;Escherichia-Shigella;Escherichia fergusonii |
| Bacteria;Proteobacteria;Gammaproteobacteria;Enterobacterales;Pasteurellaceae;Haemophilus;Haemophilus influenzae |
| Bacteria;Campylobacterota;Campylobacteria;Campylobacterales;Helicobacteraceae;Helicobacter;Helicobacter pylori |
| Bacteria;Proteobacteria;Gammaproteobacteria;Enterobacterales;Enterobacteriaceae;Klebsiella;Klebsiella aerogenes |
| Bacteria;Proteobacteria;Gammaproteobacteria;Enterobacterales;Enterobacteriaceae;Klebsiella;Klebsiella michiganensis |
| Bacteria;Proteobacteria;Gammaproteobacteria;Enterobacterales;Enterobacteriaceae;Klebsiella;Klebsiella pneumoniae |
| Bacteria;Proteobacteria;Gammaproteobacteria;Enterobacterales;Enterobacteriaceae;Klebsiella;Klebsiella quasipneumoniae |
| Bacteria;Proteobacteria;Gammaproteobacteria;Enterobacterales;Enterobacteriaceae;Klebsiella;Klebsiella variicola |
| Bacteria;Firmicutes;Bacilli;Lactobacillales;Lactobacillaceae;Lacticaseibacillus;Lacticaseibacillus paracasei |
| Bacteria;Firmicutes;Bacilli;Lactobacillales;Lactobacillaceae;Lactiplantibacillus;Lactiplantibacillus plantarum |
| Bacteria;Firmicutes;Bacilli;Lactobacillales;Streptococcaceae;Lactococcus;Lactococcus lactis |
| Bacteria;Proteobacteria;Gammaproteobacteria;Legionellales;Legionellaceae;Legionella;Legionella pneumophila |
| Bacteria;Firmicutes;Bacilli;Lactobacillales;Lactobacillaceae;Limosilactobacillus;Limosilactobacillus fermentum |
| Bacteria;Firmicutes;Bacilli;Lactobacillales;Lactobacillaceae;Limosilactobacillus;Limosilactobacillus reuteri |
| Bacteria;Firmicutes;Bacilli;Lactobacillales;Listeriaceae;Listeria;Listeria monocytogenes |

Bacteria;Proteobacteria;Gammaproteobacteria;Enterobacterales;Pasteurellaceae;Mannheimia;Mannheimia haemolytica
Bacteria;Proteobacteria;Gammaproteobacteria;Enterobacterales;Morganellaceae;Morganella;Morganella morganii
Bacteria;Actinobacteriota;Actinobacteria;Corynebacteriales;Mycobacteriaceae;Mycobacterium;Mycobacterium intracellulare
Bacteria;Actinobacteriota;Actinobacteria;Corynebacteriales;Mycobacteriaceae;Mycobacterium;Mycobacterium tuberculosis
Bacteria;Actinobacteriota;Actinobacteria;Corynebacteriales;Mycobacteriaceae;Mycobacterium;Mycobacteroides abscessus
Bacteria;Firmicutes;Bacilli;Entomoplasmatales;Entomoplasmataceae;[Mycoplasma] group;Mycoplasma mycoides
Bacteria;Firmicutes;Bacilli;Mycoplasmatales;Mycoplasmataceae;Mycoplasma;Mycoplasmopsis bovis
Bacteria;Proteobacteria;Gammaproteobacteria;Burkholderiales;Neisseriaceae;Neisseria;Neisseria gonorrhoeae
Bacteria;Proteobacteria;Gammaproteobacteria;Burkholderiales;Neisseriaceae;Neisseria;Neisseria meningitidis
Bacteria;Proteobacteria;Gammaproteobacteria;Enterobacterales;Pasteurellaceae;Pasteurella;Pasteurella multocida
Bacteria;Proteobacteria;Gammaproteobacteria;Piscirickettsiales;Piscirickettsiaceae;Piscirickettsia;Piscirickettsia salmonis
Bacteria;Proteobacteria;Gammaproteobacteria;Enterobacterales;Morganellaceae;Proteus;Proteus mirabilis
Bacteria;Proteobacteria;Gammaproteobacteria;Pseudomonadales;Pseudomonadaceae;Pseudomonas;Pseudomonas aeruginosa
Bacteria;Proteobacteria;Gammaproteobacteria;Pseudomonadales;Pseudomonadaceae;Pseudomonas;Pseudomonas chlororaphis
Bacteria;Proteobacteria;Gammaproteobacteria;Burkholderiales;Burkholderiaceae;Ralstonia;Ralstonia solanacearum
Bacteria;Proteobacteria;Gammaproteobacteria;Enterobacterales;Enterobacteriaceae;Salmonella;Salmonella enterica
Bacteria;Proteobacteria;Gammaproteobacteria;Enterobacterales;Yersiniaceae;Serratia;Serratia marcescens
Bacteria;Firmicutes;Bacilli;Staphylococcales;Staphylococcaceae;Staphylococcus;Staphylococcus aureus
Bacteria;Firmicutes;Bacilli;Staphylococcales;Staphylococcaceae;Staphylococcus;Staphylococcus epidermidis
Bacteria;Firmicutes;Bacilli;Staphylococcales;Staphylococcaceae;Staphylococcus;Staphylococcus pseudintermedius
Bacteria;Firmicutes;Bacilli;Lactobacillales;Streptococcaceae;Streptococcus;Streptococcus agalactiae
Bacteria;Firmicutes;Bacilli;Lactobacillales;Streptococcaceae;Streptococcus;Streptococcus dysgalactiae
Bacteria;Firmicutes;Bacilli;Lactobacillales;Streptococcaceae;Streptococcus;Streptococcus pneumoniae
Bacteria;Firmicutes;Bacilli;Lactobacillales;Streptococcaceae;Streptococcus;Streptococcus pyogenes
Bacteria;Firmicutes;Bacilli;Lactobacillales;Streptococcaceae;Streptococcus;Streptococcus suis
Bacteria;Firmicutes;Bacilli;Lactobacillales;Streptococcaceae;Streptococcus;Streptococcus thermophilus
Bacteria;Proteobacteria;Gammaproteobacteria;Enterobacterales;Vibrionaceae;Vibrio;Vibrio cholerae
Bacteria;Proteobacteria;Gammaproteobacteria;Enterobacterales;Vibrionaceae;Vibrio;Vibrio parahaemolyticus
Bacteria;Proteobacteria;Gammaproteobacteria;Xanthomonadales;Xanthomonadaceae;Xanthomonas;Xanthomonas citri
Bacteria;Proteobacteria;Gammaproteobacteria;Xanthomonadales;Xanthomonadaceae;Xanthomonas;Xanthomonas oryzae
Bacteria;Proteobacteria;Gammaproteobacteria;Xanthomonadales;Xanthomonadaceae;Xylella;Xylella fastidiosa
Bacteria;Proteobacteria;Gammaproteobacteria;Enterobacterales;Yersiniaceae;Yersinia;Yersinia pestis

**Table S2. GLMMs of palindrome avoidance for (a) k=4 and (b) 6 across the genomic dataset, controlling for phylogeny.** Models are fitted to the mean exceptionality score for each species. Species with fewer than 3 isolates are excluded.

**(a) Palindrome avoidance modelling (K=4)**

| | | | Variance explained (%) | Mean estimate | Lower (95% CI) | Upper (95%) | Effective sample size | pMCMC |
|---|---|---|---|---|---|---|---|---|
| Subsampling (bp) | 10000 | Intercept | - | -0.71 | -1.34 | -0.07 | 1000.00 | 0.04 |
| Genomes | 4545 | Category (relative to core): Non-core | 0.25 | -0.09 | -0.21 | 0.02 | 1000.00 | 0.14 |
| Species | 54 | Category (relative to core): Plasmid-borne | 0.00 | 0.01 | -0.10 | 0.11 | 894.41 | 0.88 |
| | | Phylogeny | 84.11 | | | | | |
| | | No. of genomes | 2.43 | | | | | |
| | | Residuals | 13.21 | | | | | |

| | | | Variance explained (%) | Mean estimate | Lower (95% CI) | Upper (95%) | Effective sample size | pMCMC |
|---|---|---|---|---|---|---|---|---|
| Subsampling (bp) | 50000 | Intercept | - | -1.27 | -2.43 | 0.34 | 1000.00 | 0.07 |
| Genomes | 3905 | Category (relative to core): Non-core | 0.12 | -0.11 | -0.33 | 0.08 | 1000.00 | 0.26 |
| Species | 49 | Category (relative to core): Plasmid-borne | 0.03 | 0.05 | -0.15 | 0.24 | 1000.00 | 0.59 |
| | | Phylogeny | 88.09 | | | | | |
| | | No. of genomes | 1.22 | | | | | |
| | | Residuals | 10.54 | | | | | |

| | | | Variance explained (%) | Mean estimate | Lower (95% CI) | Upper (95%) | Effective sample size | pMCMC |
|---|---|---|---|---|---|---|---|---|
| Subsampling (bp) | 100000 | Intercept | - | -1.51 | -3.38 | 0.66 | 1000.00 | 0.14 |
| Genomes | 3090 | Category (relative to core): Non-core | 0.35 | -0.28 | -0.58 | 0.01 | 1103.41 | 0.05 |
| Species | 42 | Category (relative to core): Plasmid-borne | 0.01 | -0.05 | -0.33 | 0.24 | 1000.00 | 0.73 |
| | | Phylogeny | 89.49 | | | | | |
| | | No. of genomes | 0.96 | | | | | |
| | | Residuals | 9.20 | | | | | |

**(b) Palindrome avoidance modelling (K=6)**

| Variance explained (%) | Mean estimate | Lower (95% CI) | Upper (95%) | Effective sample size | pMCMC |
|---|---|---|---|---|---|

| | | | | | | | | |
|---|---|---|---|---|---|---|---|---|
| Subsampling (bp) | 10000 | Intercept | - | -0.17 | -0.35 | 0.02 | 1000.00 | 0.07 |
| Genomes | 4545 | Category (relative to core): Non-core | 0.01 | 0.00 | -0.04 | 0.05 | 1130.69 | 0.86 |
| Species | 54 | Category (relative to core): Plasmid-borne | 3.89 | -0.10 | -0.15 | -0.05 | 1000.00 | 0.00 |
| | | Phylogeny | 66.00 | | | | | |
| | | No. of genomes | 3.39 | | | | | |
| | | Residuals | 26.72 | | | | | |
| | | | Variance explained (%) | Mean estimate | Lower (95% CI) | Upper (95%) | Effective sample size | pMCMC |
| Subsampling (bp) | 50000 | Intercept | - | -0.45 | -0.65 | -0.25 | 1000.00 | 0.00 |
| Genomes | 3905 | Category (relative to core): Non-core | 0.05 | 0.01 | -0.06 | 0.08 | 1095.37 | 0.67 |
| Species | 49 | Category (relative to core): Plasmid-borne | 7.10 | -0.16 | -0.22 | -0.09 | 836.92 | 0.00 |
| | | Phylogeny | 49.74 | | | | | |
| | | No. of genomes | 6.28 | | | | | |
| | | Residuals | 36.83 | | | | | |
| | | | Variance explained (%) | Mean estimate | Lower (95% CI) | Upper (95%) | Effective sample size | pMCMC |
| Subsampling (bp) | 100000 | Intercept | - | -0.56 | -0.85 | -0.30 | 1000.00 | 0.00 |
| Genomes | 3090 | Category (relative to core): Non-core | 0.03 | -0.01 | -0.10 | 0.07 | 1060.04 | 0.74 |
| Species | 42 | Category (relative to core): Plasmid-borne | 4.85 | -0.17 | -0.27 | -0.09 | 1000.00 | 0.00 |
| | | Phylogeny | 59.16 | | | | | |
| | | No. of genomes | 2.84 | | | | | |
| | | Residuals | 33.12 | | | | | |

**Table S3. Modelling results for orphan MTase carriage as a function of PTU size and host range.** Size dominates as a factor for all PTUs, but when only considering large PTUs host range explains a significant amount of variance. This modelling only considers MTases which are not part of a putative R-M system on the plasmid.

(a) All PTUs

| | Df | Sum Sq | Mean Sq | F value | Pr(>F) | Variance explained (%) |
|---|---|---|---|---|---|---|
| log10(size) | 1 | 1.77 | 1.77 | 33.75 | <0.01 | 10.88 |
| Host range | 1 | 0.19 | 0.19 | 3.54 | 0.06 | 1.14 |
| Residuals | 273 | 14.32 | 0.05 | NA | NA | 87.98 |

(b) PTUs >100kb

| | Df | Sum Sq | Mean Sq | F value | Pr(>F) | Variance explained (%) |
|---|---|---|---|---|---|---|
| log10(size) | 1 | 0.01 | 0.01 | 0.12 | 0.73 | 0.14 |
| Host range | 1 | 2.15 | 2.15 | 22.24 | <0.01 | 27.34 |
| Residuals | 59 | 5.70 | 0.10 | NA | NA | 72.52 |

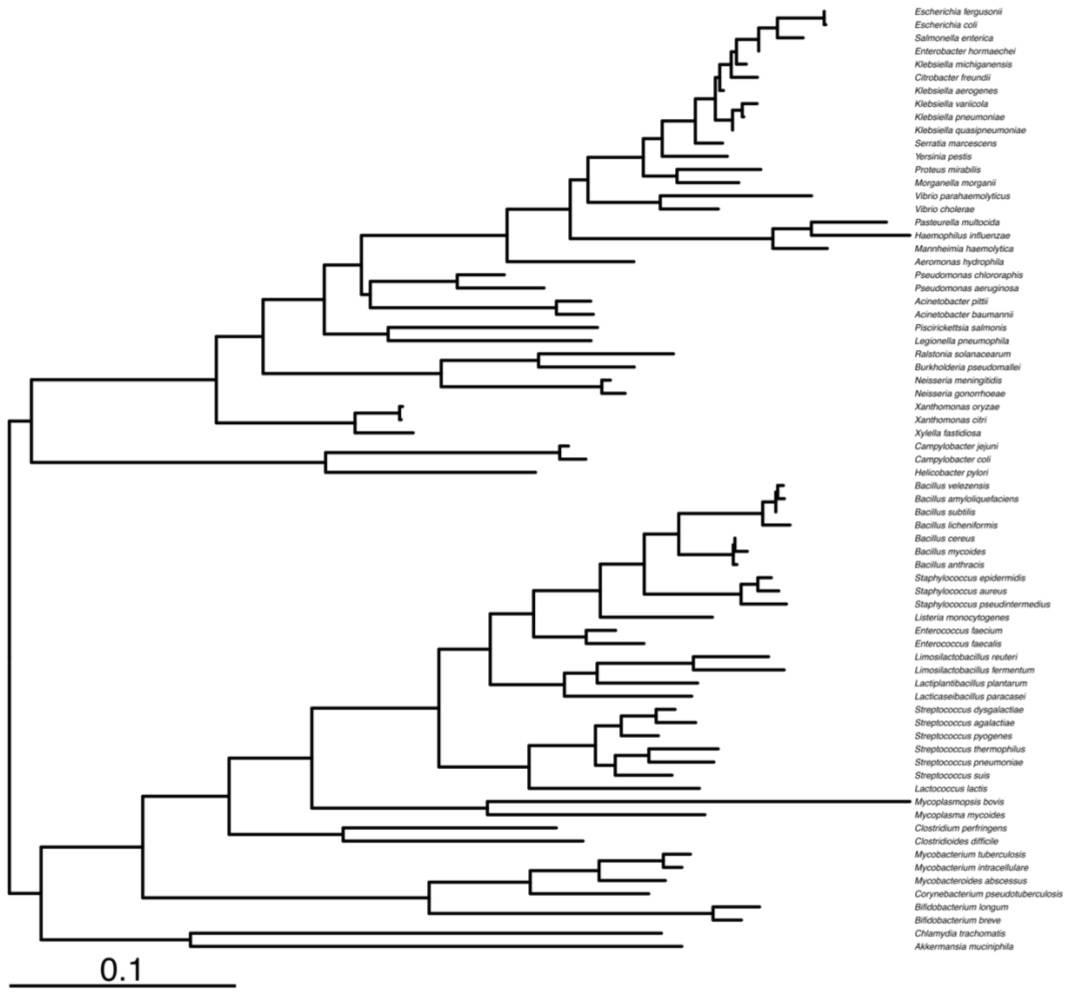


**Fig. S1. Midpoint-rooted 16S rRNA gene tree of the host species in the genomic dataset.** This phylogeny was produced from a multiple sequence alignment with FastTree (see Methods) and was used to control for phylogenetic structure in modelling analysis. Branch lengths represent average substitutions per site, with scale bar representing 0.1 substitutions per site.

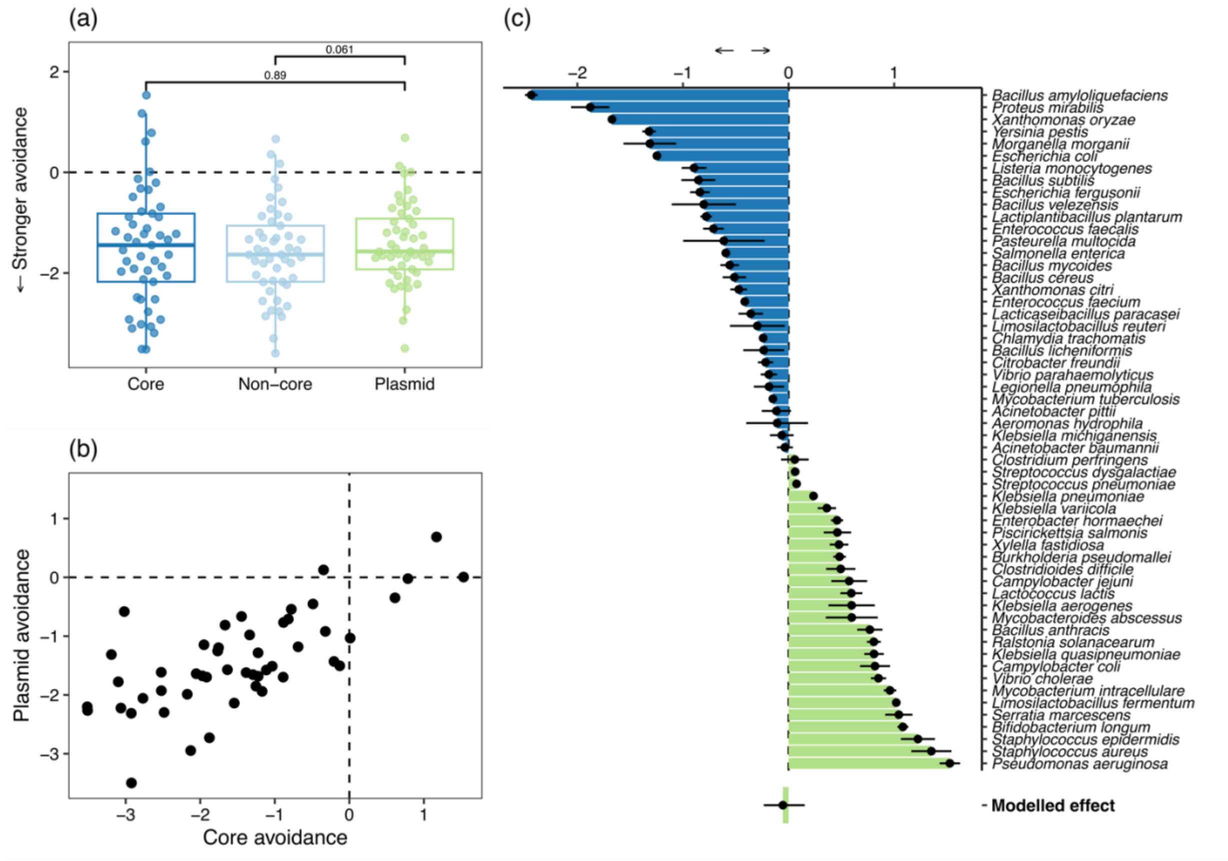


**Fig. S2. Palindrome avoidance for k=4.** (a) 4-bp palindrome avoidance scores do not show differences between plasmid-borne genes compared to core and non-core chromosomal genes ($p>0.05$, two-sided Wilcoxon paired test). (b) Mean avoidance is strongly structured by species, with a strong correlation between avoidance in core and plasmid-borne genes. (c) Relative palindrome avoidance for species for core vs. plasmid-borne genes (>0 denotes greater avoidance in plasmid-borne genes). Points are mean, error bars show standard error. The modelled effect is from a phylogenetically-controlled GLMM (see Methods). Data shown are mean avoidance scores of 4-bp palindromes (12 possible palindromes) calculated with R'MES after pangenome construction then subsampling each per-isolate pangenome component to 50kbp i.e. only genomes with at least 50kbp are included (3,912 isolate genomes across 44 species). Only species with at least 3 genomes meeting these criteria are shown. Subsampling to other fixed lengths for pangenome components gives very similar results.

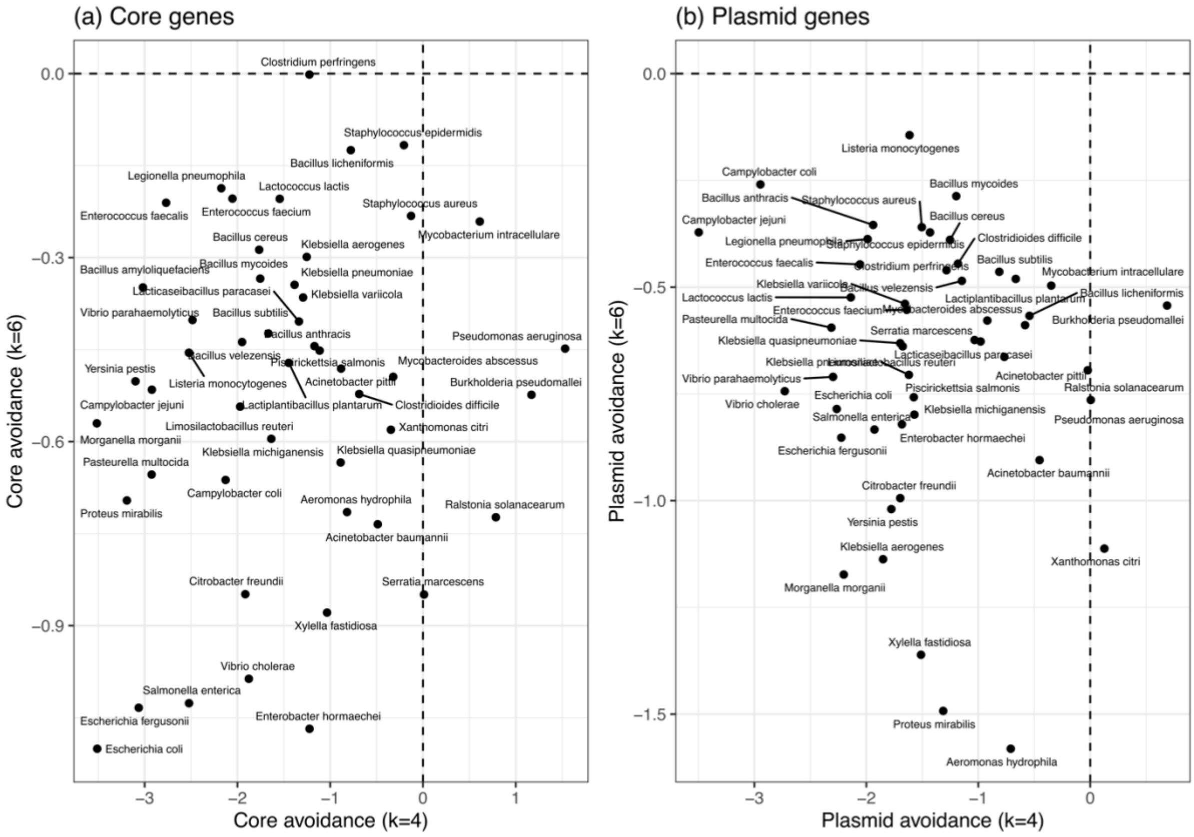


**Fig. S3. Lack of correlation between palindrome avoidance for k=4 and k=6.** Mean avoidance of palindromes of length k=4 and k=6 is not correlated across bacterial species for (a) core or (b) plasmid gene

k=4

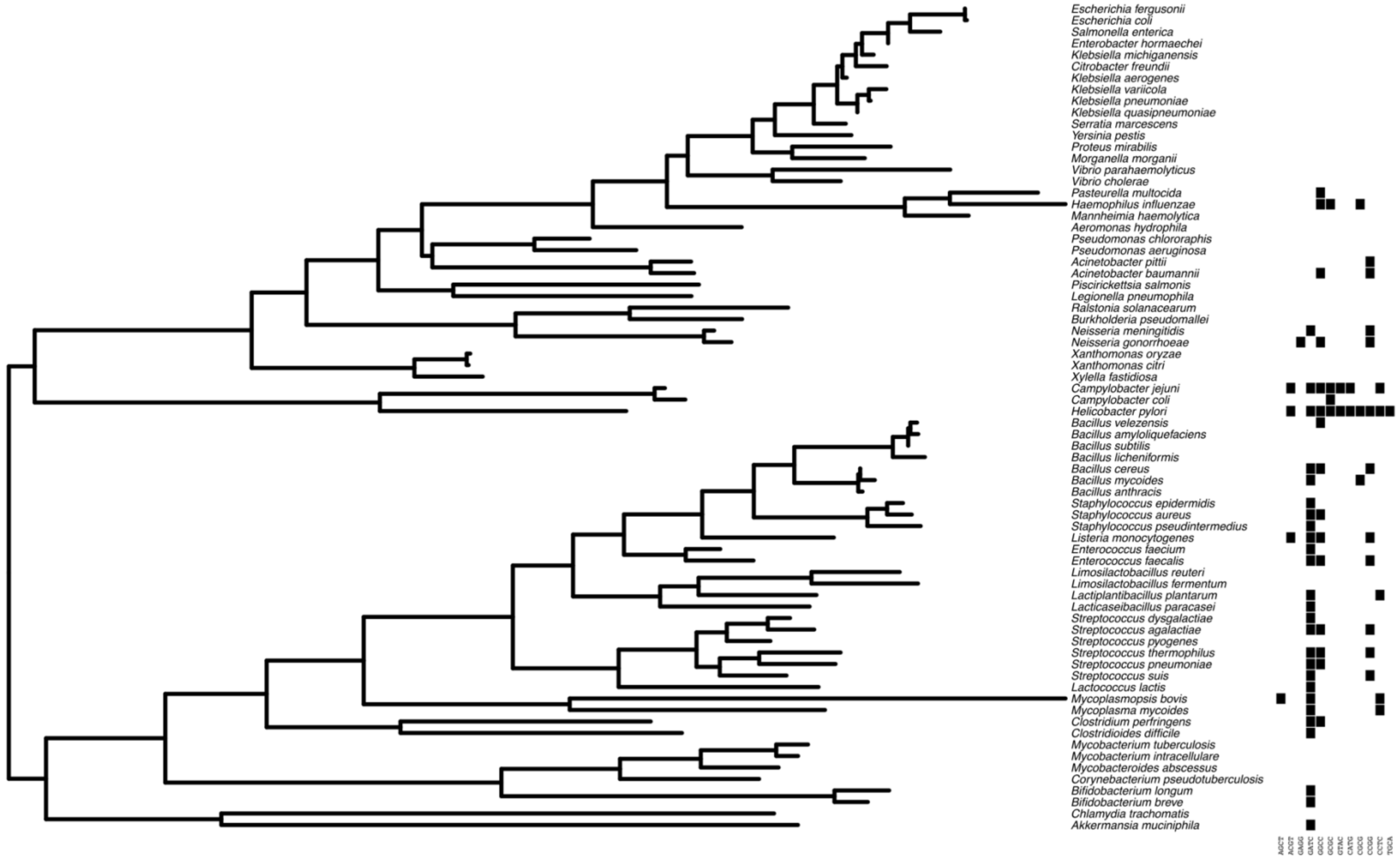


**Fig. S4. Heatmaps of the presence of Type II R-M systems targeting targets of length k=4.** Phylogenetic tree as in Fig. S1.

k=5

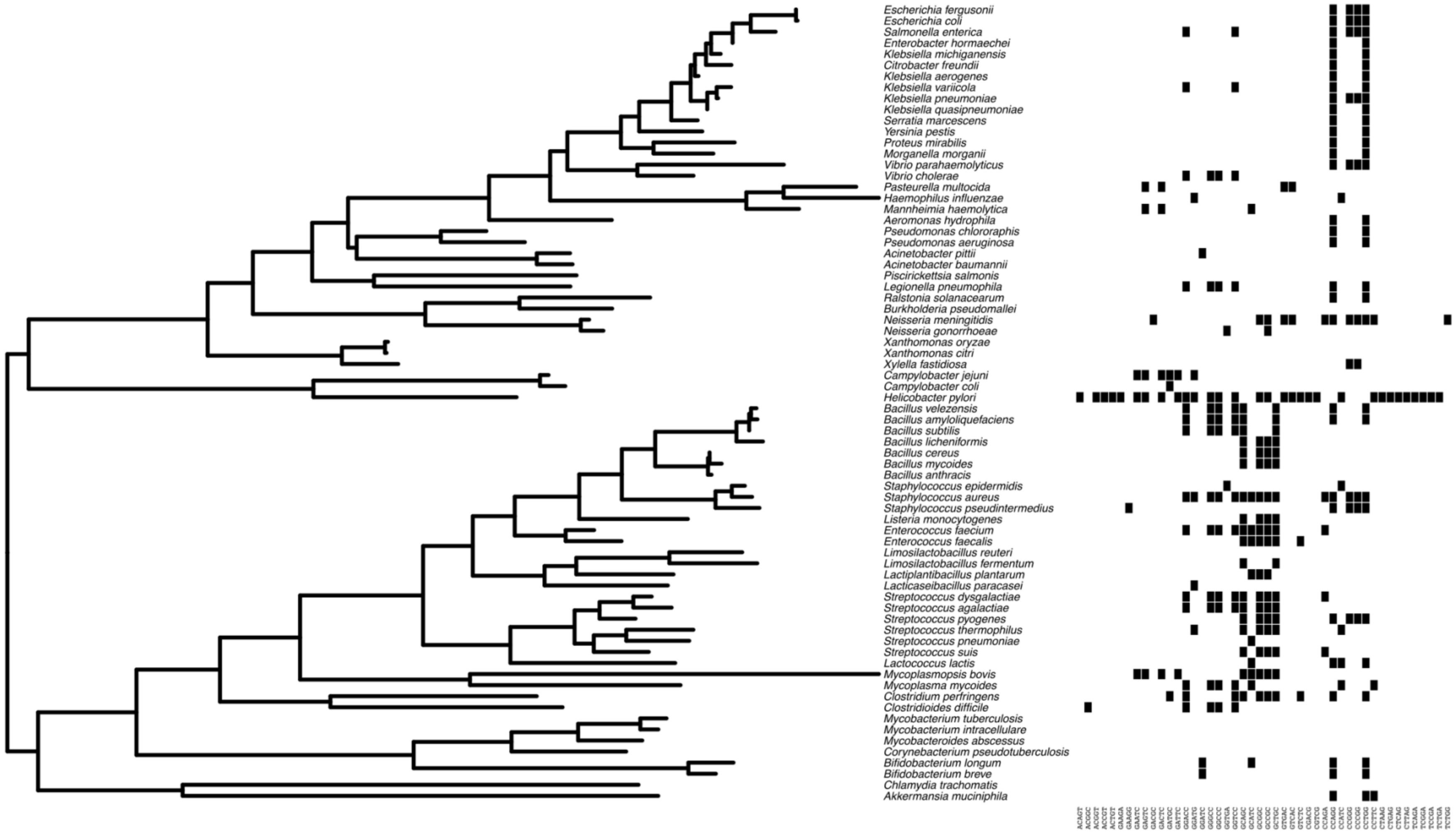


**Fig. S5. Heatmaps of the presence of Type II R-M systems targeting targets of length k=5.** Phylogenetic tree is from Fig. S3.

k=6

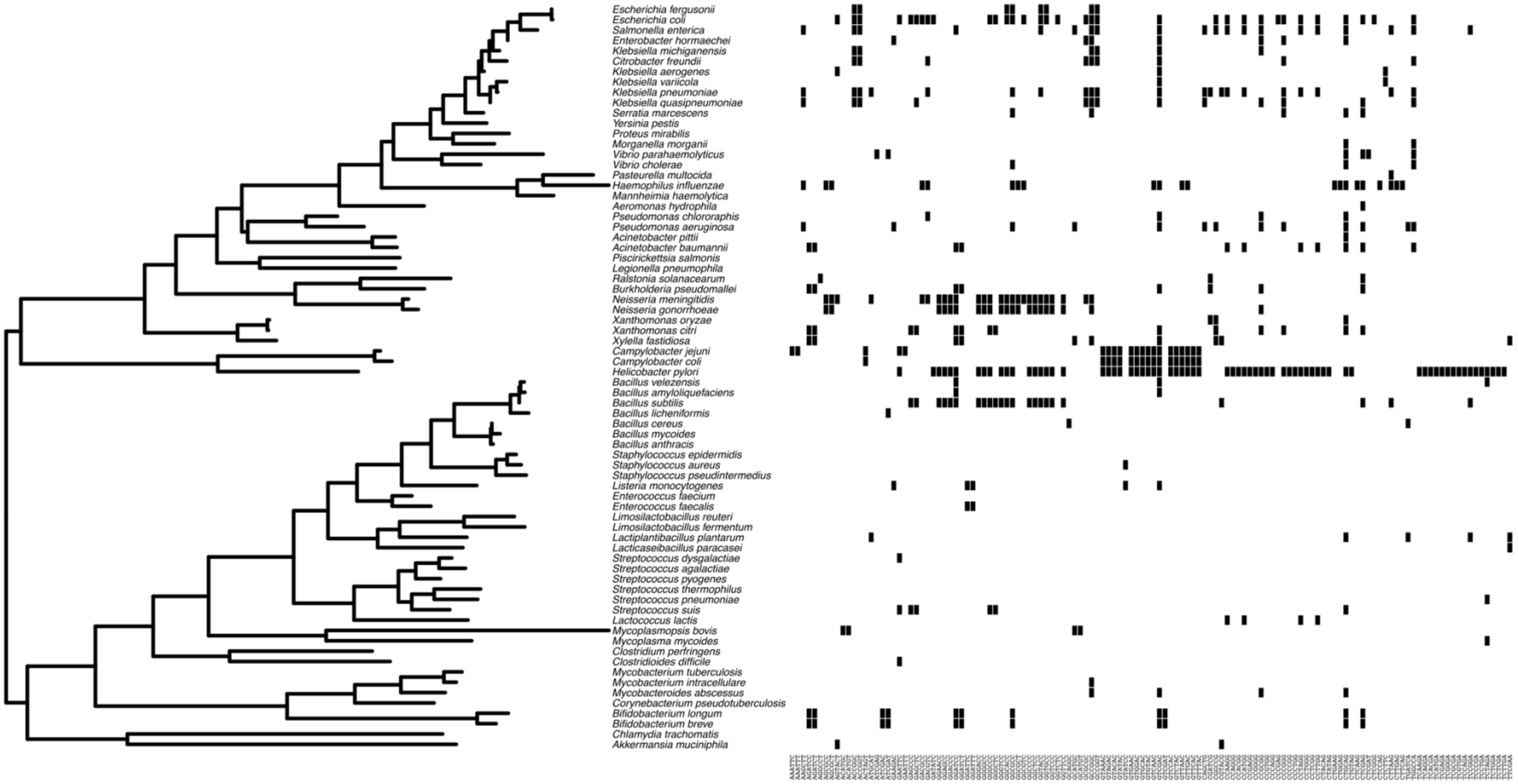


**Fig. S6. Heatmaps of the presence of Type II R-M systems targeting targets of length k=6.** Phylogenetic tree is from Fig. S3.

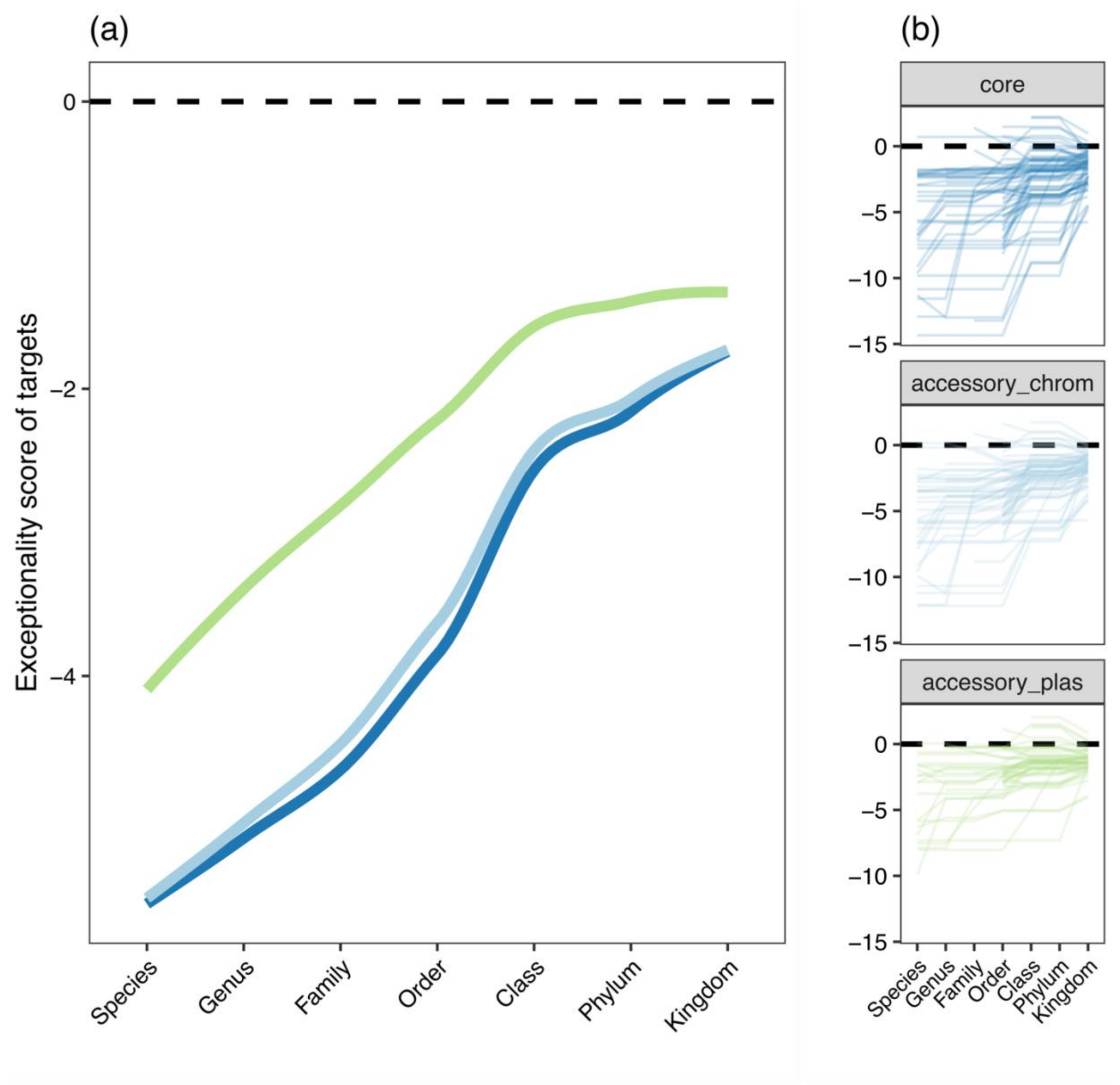


**Fig. S7. The taxonomic distribution of R-M systems correlates with avoidance of their targets in the pangenome for 4-mer targets.** (a) The total unique 4-mer targets recognised by Type II R-M systems detected at least once in each species. Phylogeny adapted from Zhu et al. ( 2019). (b) Exceptionality scores for 4-mers by pangenome component as a function of the taxonomic hierarchy of R-M targets, averaged over all species. Negative scores are associated with avoidance of targets. Mean avoidance scores are shown for core chromosomal genes (dark blue), non-core chromosomal genes (light blue) and plasmid-borne genes (green). Levels on the x-axis for the taxonomic hierarchy of targets are inclusive i.e. genus includes targets of R-M systems within-species (see Fig. 2). Subsampling is to 50kbp for each within-isolate pangenome component. Inset panels on right show results for individual species.

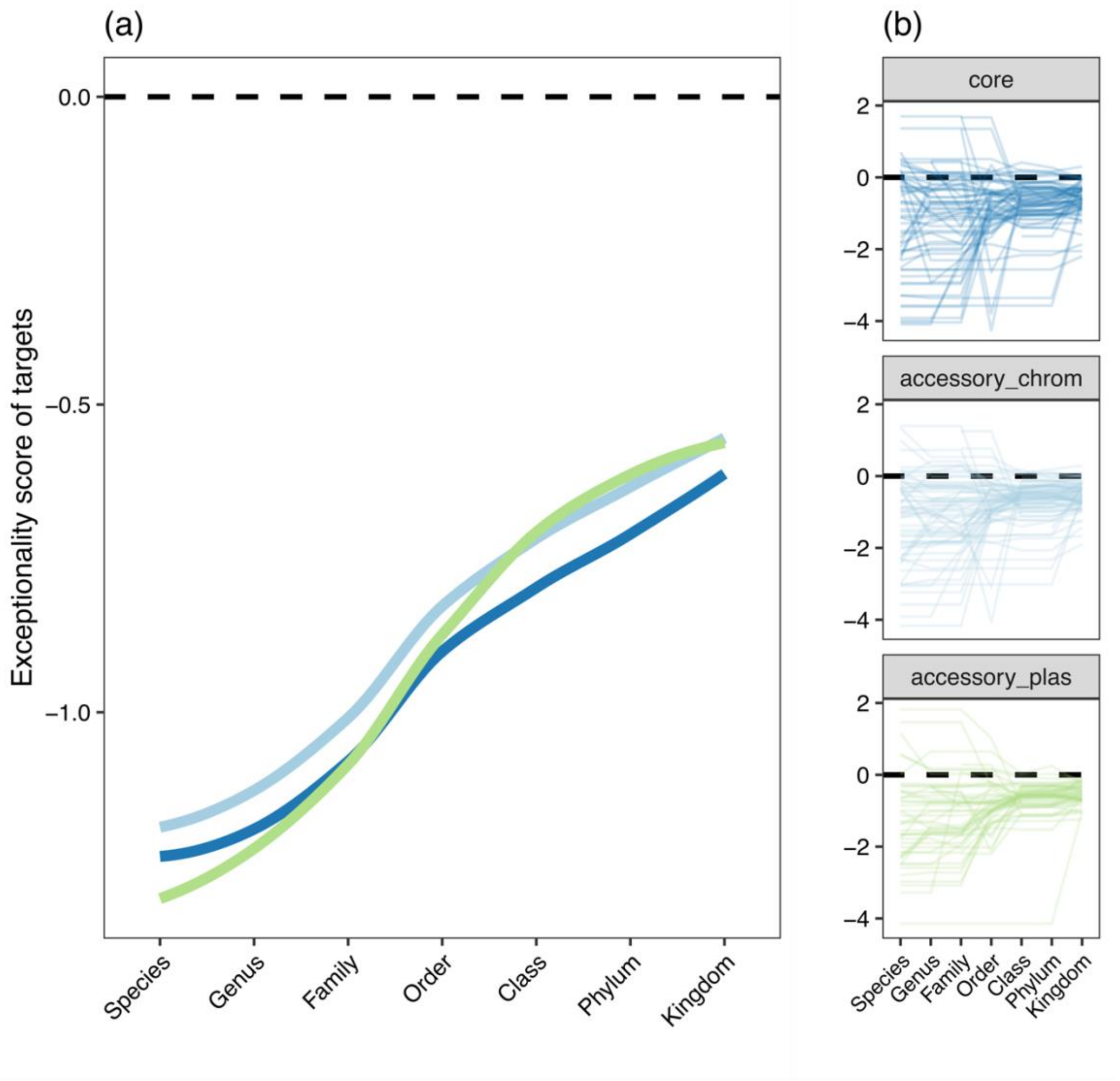


**Fig. S8. The taxonomic distribution of R-M systems correlates with avoidance of their targets in the pangenome for 5-mer targets.** (a) The total unique 5-mer targets recognised by Type II R-M systems detected at least once in each species. Phylogeny adapted from Zhu et al. ( 2019). (b) Exceptionality scores for 5-mers by pangenome component as a function of the taxonomic hierarchy of R-M targets, averaged over all species. Negative scores are associated with avoidance of targets. Mean avoidance scores are shown for core chromosomal genes (dark blue), non-core chromosomal genes (light blue) and plasmid-borne genes (green). Levels on the x-axis for the taxonomic hierarchy of targets are inclusive i.e. genus includes targets of R-M systems within-species (see Fig. 2). Subsampling is to 50kbp for each within-isolate pangenome component. Inset panels on right show results for individual species.

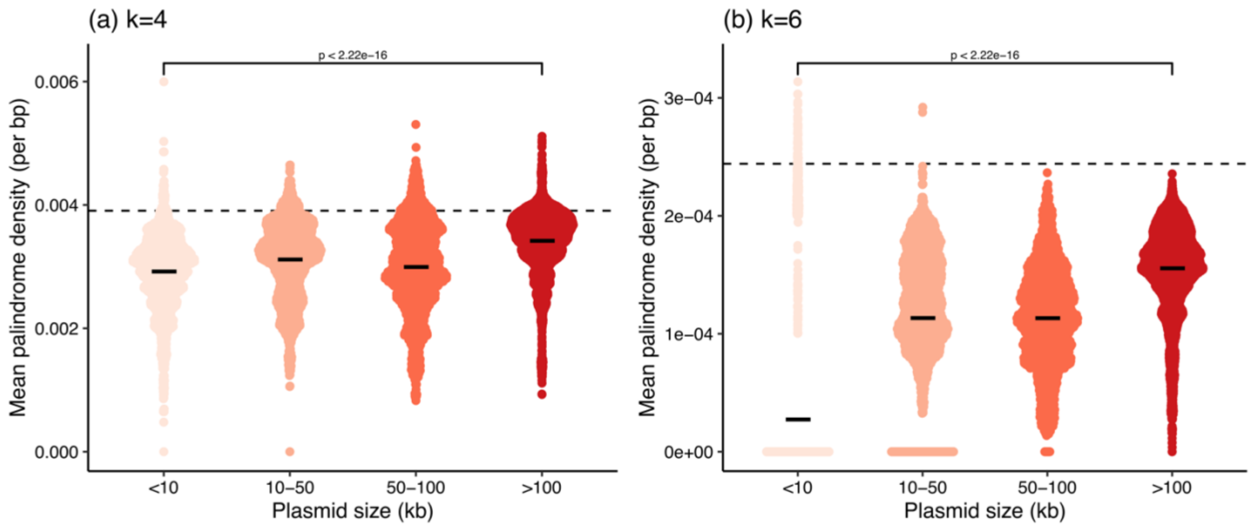


**Fig. S9. Palindrome densities scale with plasmid size.**
Similar to Fig. 3, but for the mean densities of palindromes of (a) k=4 and (b) k=6. Each point is a plasmid (or secondary chromosome for a small number of species, see Methods) in a genome.

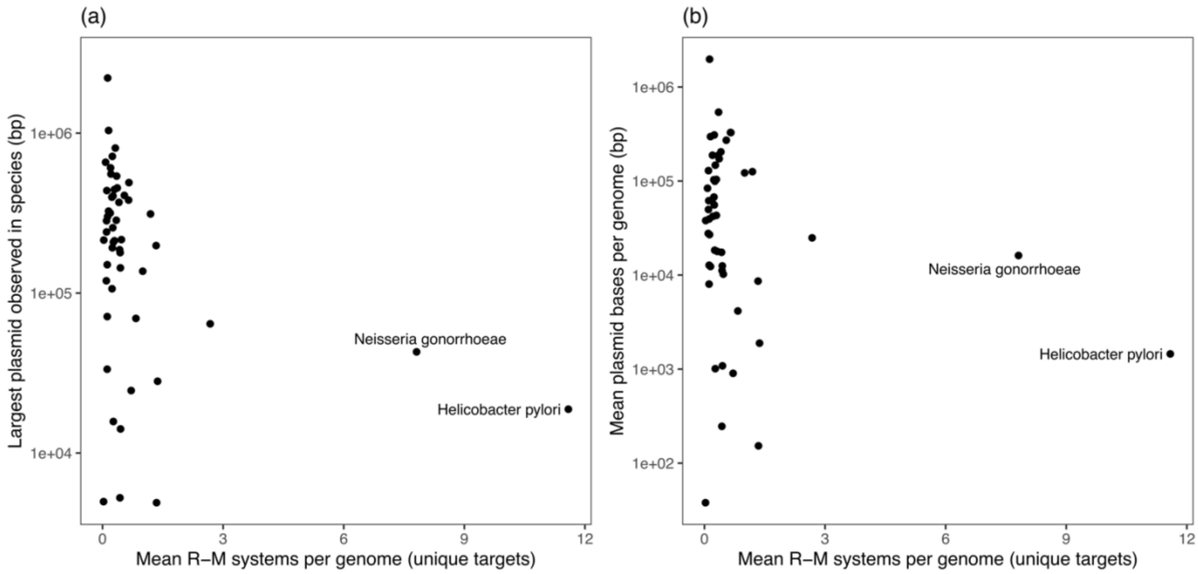


**Fig. S10. Species with many and diverse R-M systems do not have large plasmids.**
(a) Largest observed plasmid and (b) mean plasmid bases per genome for n=57 species with both R-M systems and plasmids. Though there is not a linear relationship, it is clear that the two outlier 'R-M-rich' species do not have large plasmids, consistent with the idea that a large number of diverse R-M systems in a genome would exert non-overlapping selective pressures that should exclude the possibility of large plasmids persisting.

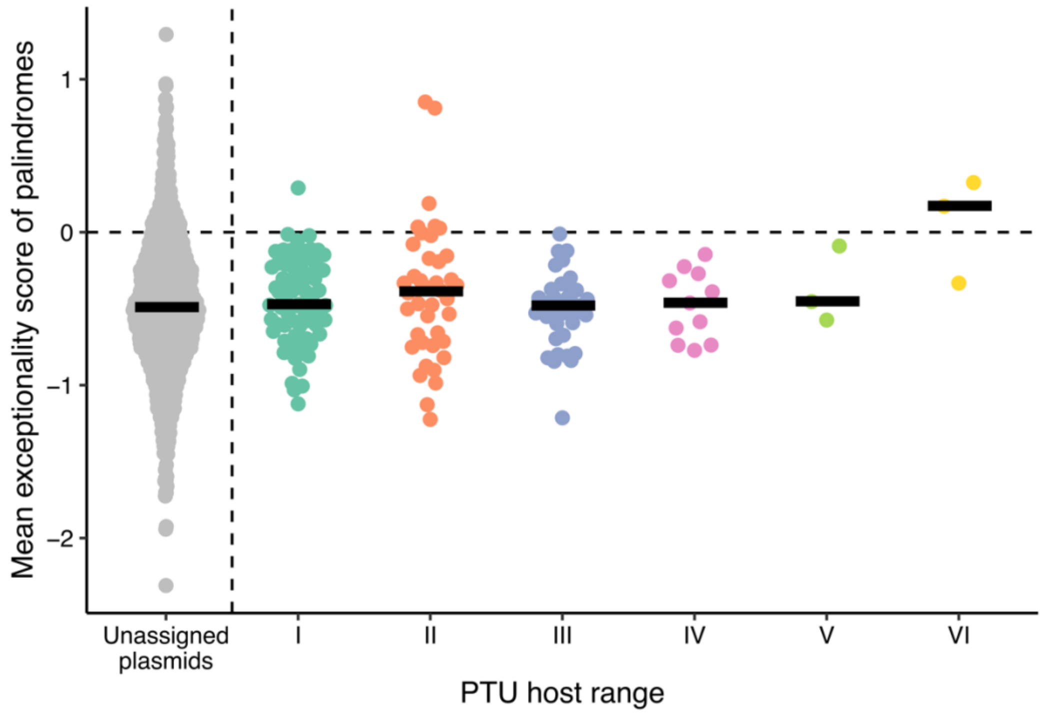


**Fig. S11. PTU host range is not associated with greater avoidance of 4-bp palindromes.** As Fig. 4, but for 4-bp palindromes. Each point is one PTU (mean exceptionality score) apart from unassigned plasmids (those not classified into a PTU) and lines show median within host ranges.

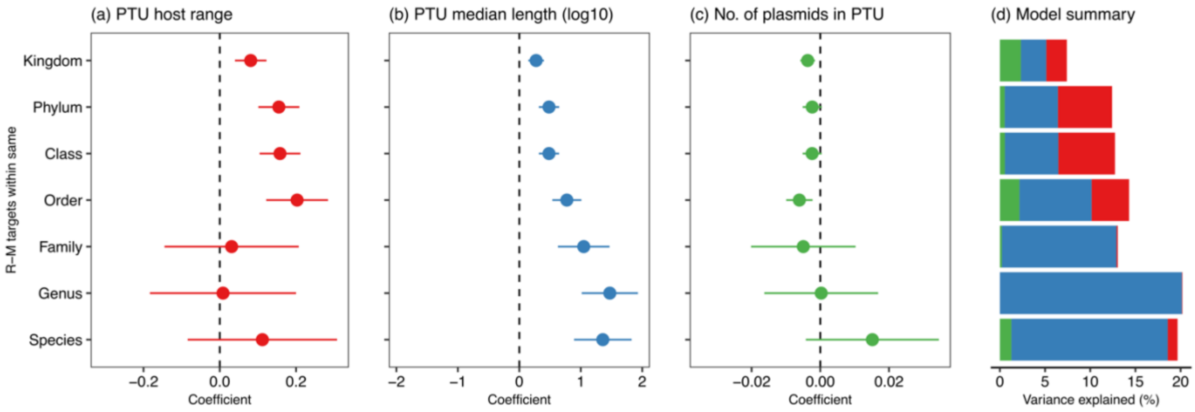


**Fig. S12. Small PTUs have stronger avoidance of R-M targets of length k=4.**
(a-c) Coefficients in linear models (mean estimates with standard error shown by errorbars) for the exceptionality score of R-M targets. A different model was run for each possible of level of R-M targets within the taxonomic hierarchy, from R-M targets of R-M systems within-species to within-kingdom, with three variables for each PTU: host range, median length, and number of plasmids. (d) Total variance explained by models.

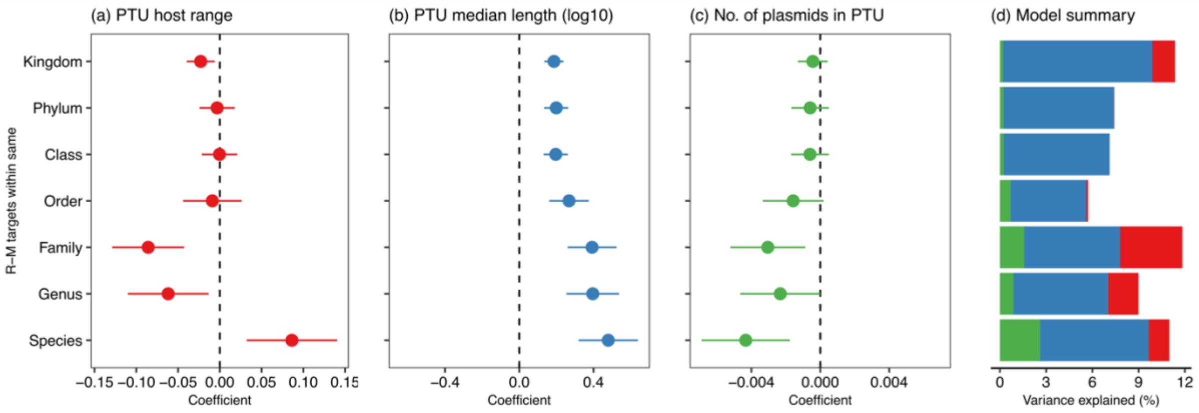


**Fig. S13. Small PTUs have stronger avoidance of R-M targets of length k=5.**
(a-c) Coefficients in linear models (mean estimates with standard error shown by errorbars) for the exceptionality score of R-M targets. A different model was run for each possible of level of R-M targets within the taxonomic hierarchy, from R-M targets of R-M systems within-species to within-kingdom, with three variables for each PTU: host range, median length, and number of plasmids. (d) Total variance explained by models.

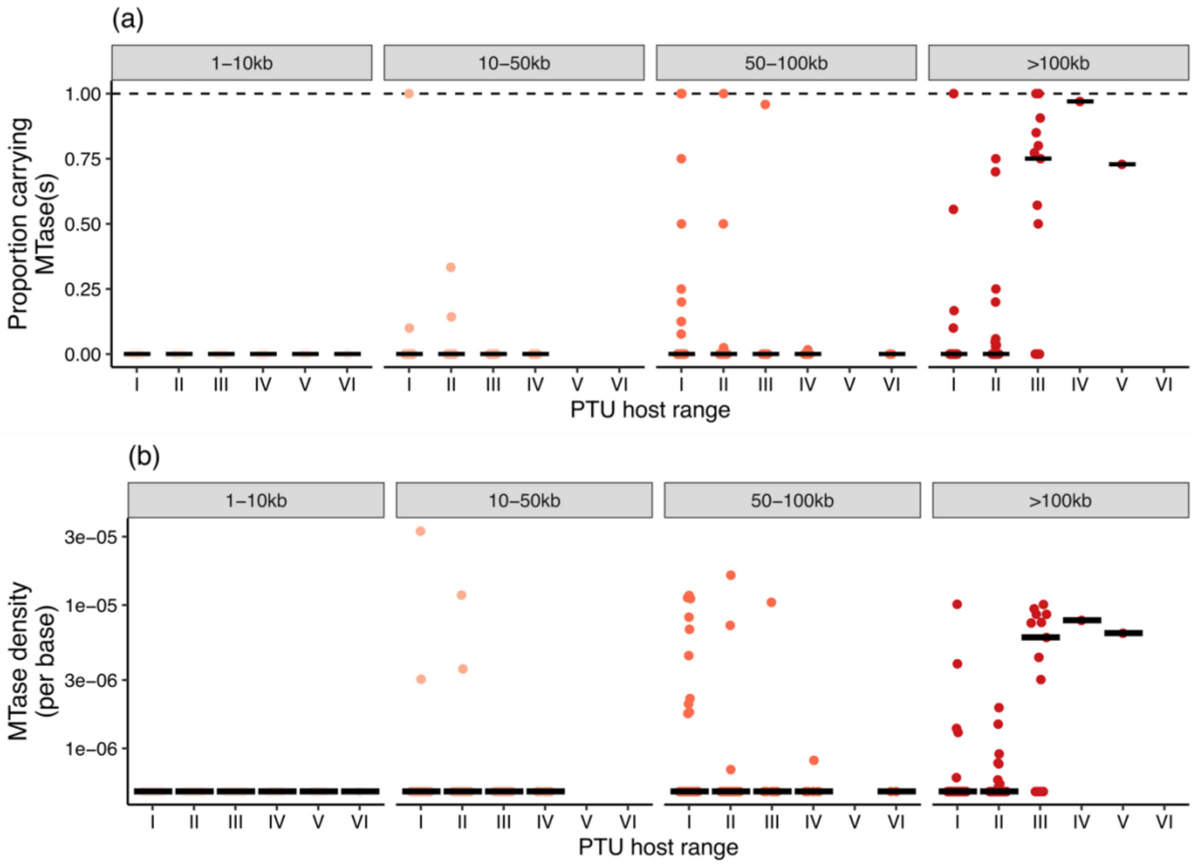


**Fig. S14. Large plasmids with a broad host range carry orphan MTases.** Each point represents one plasmid taxonomic unit (PTU), n=271 total. Plots are faceted by PTU size. (a) Proportion of plasmids within a PTU carrying at least one MTase. (b) Mean density of MTases (total number divided by median size of PTU). Boxplots show median.
